# SimSpace: a comprehensive in-silico spatial omics data simulation and modeling framework

**DOI:** 10.1101/2025.07.18.665587

**Authors:** Tianxiao Zhao, Katherine Zhang, Michelle Hollenberg, Wen Zhou, David Fenyö

**Affiliations:** Institute for Systems Genetics, NYU Grossman School of Medicine, New York, NY, USA; Department of Biochemistry and Molecular Pharmacology, NYU Grossman School of Medicine, New York, NY, USA; Department of Biostatistics, NYU School of Global Public Health, New York, NY, USA

## Abstract

Spatial omics technologies provide rich measurements of molecular states in their native tissue context, but the development and benchmarking of computational methods remain constrained by limited access to datasets with known ground truth. We present *SimSpace*, a flexible framework for simulating spatial omics data for method development, stress-testing, and controlled in silico experiments. SimSpace uses a hierarchical generative model that couples tissue-scale spatial organization, local cell-type interactions, and modular omics-profile generation. Its Markov random field-based spatial model enables controllable simulation of niche structure, cell-type co-localization, phenotype-dependent density, and spatially mediated molecular interactions. SimSpace supports both reference-free simulations under user-defined generative assumptions and reference-guided simulations that calibrate spatial parameters to real datasets while generating new spatial realizations. Across Xenium, MERFISH, and CODEX examples, SimSpace reproduced multiple evaluated spatial statistics, preserved key domain-level structure in reference-guided settings, and supported benchmarking of cell-type deconvolution and spatially variable gene detection methods. We further show that inferred interaction parameters can be perturbed to generate interpretable counterfactual spatial proteomics scenarios. SimSpace is implemented as an open-source Python package with reproducible workflows for rigorous benchmarking and methodological development in spatial omics.

## Introduction

Spatial omics technologies are rapidly transforming the understanding of how tissue function emerges from the spatial organization of cells. By profiling molecular features in their native tissue context, spatial transcriptomics and related spatial omics technologies have provided new views of tissue architecture, cellular neighborhoods, and intercellular communication. These spatial patterns are central to many biological processes, including tissue development, embryogenesis, neuroscience, tumor evolution, and immune organization [1–4].

This technological progress has motivated a rapidly growing ecosystem of computational methods for spatial omics analysis [5, 6]. These methods address a wide range of tasks, including cell-type deconvolution [7–11], spatially variable gene detection [12–17], spatial imputation [18, 19], and spatial domain recognition [20–22]. Despite this progress, rigorous benchmarking remains difficult because real datasets rarely provide complete ground truth for spatial cell organization, molecular variation, and cell–cell interaction structure.

In silico simulation offers a promising complementary strategy for benchmarking spatial omics methods by generating synthetic datasets with known ground truth. However, the value of a simulator depends critically on whether it can reproduce the statistical and biological structure that downstream methods are designed to detect. Many simulation settings used in methodological studies are intentionally simple [23–26], whereas others rely on assumptions tailored to a specific methodological target [14, 27, 28]. Such designs are useful for controlled validation, but they may lead to incomplete or biased assessments when used as general benchmarks for spatial omics methods. A key unmet need is a flexible simulator that can generate biologically plausible tissue architectures while retaining explicit, interpretable ground-truth mechanisms. Existing simulation tools can be broadly categorized as “reference-based” or “reference-free”. Reference-based approaches condition on an observed spatial omics dataset and generate new molecular profiles using information learned from that reference. Examples include scDesign3 [29], SRTsim [30], and scCube [31]. These approaches can produce realistic molecular distributions, but current implementations often reuse the original spatial coordinate scaffold or assign simulated profiles back to reference-derived tissue locations. As a result, the simulated data inherit much of the coordinate geometry and spatial organization of the input tissue, which limits their ability to generate new tissue shapes, alternative niche arrangements, or controlled perturbations of spatial cell–cell interactions.

Reference-free approaches, in contrast, generate spatial coordinates and tissue structure de novo without requiring an observed spatial reference. SRTsim and scCube provide reference-free workflows based on pre-defined geometric templates, user-specified layouts, or random spatial sampling, whereas scMultiSim [32] introduces low-dimensional spatial structure through basis functions or grid-based gene regulatory network processes. These methods are effective for generating coarse spatial variation, but they often treat coordinates, cell labels, and molecular profiles as partially separate components. Therefore, they may not explicitly represent the hierarchical organization of real tissues, where tissue domains, local cell-type composition, cellular proximity, and molecular interaction programs are coupled across spatial scales.

This distinction highlights an important statistical and computational challenge. Spatial omics simulators should not only generate visually plausible coordinates or realistic feature distributions; they should also specify how tissue-level domains, cell-type neighborhoods, and molecular measurements jointly arise from a common generative mechanism. Such a model-based formulation is essential for benchmarking because it makes the relevant sources of ground truth explicit, including domain structure, local spatial autocorrelation, cell-type co-localization, phenotype-dependent density, marker-feature effects, and spatially mediated molecular interactions. At the same time, different simulation settings serve complementary purposes. Reference-free simulations cannot capture the full anatomical complexity of real tissues, but they provide controlled generative models for evaluating whether computational methods behave as expected. Reference-based simulations, in contrast, leverage biological structure from real data and can generate synthetic tissues that preserve key spatial features while still providing known ground truth. These complementary roles motivate a flexible framework that supports both controlled reference-free experiments and reference-guided simulation calibrated to real tissues.

To address these needs, we present SimSpace, a simulation and benchmarking framework for generating synthetic spatial omics datasets with controllable tissue architecture and modular molecular profiles. SimSpace is built on a hierarchical generative model (**Fig. 1A**) that first generates spatial cell locations and domains, then assigns cell phenotypes conditional on spatial context, and finally simulates omics profiles conditional on both phenotype and location. A central methodological feature of SimSpace is its two-level spatial dependence model. At the tissue scale, SimSpace uses a niche-level Markov random field (MRF) to generate spatially coherent domains. At the cellular scale, it uses a second, niche-conditioned MRF to generate local cell-type organization within and across these domains. This hierarchical affinity structure allows users to control both niche–niche organization and cell-type–cell-type proximity, providing interpretable ground truth for spatial autocorrelation, co-localization, and cell–cell interaction patterns.

**Figure 1.**
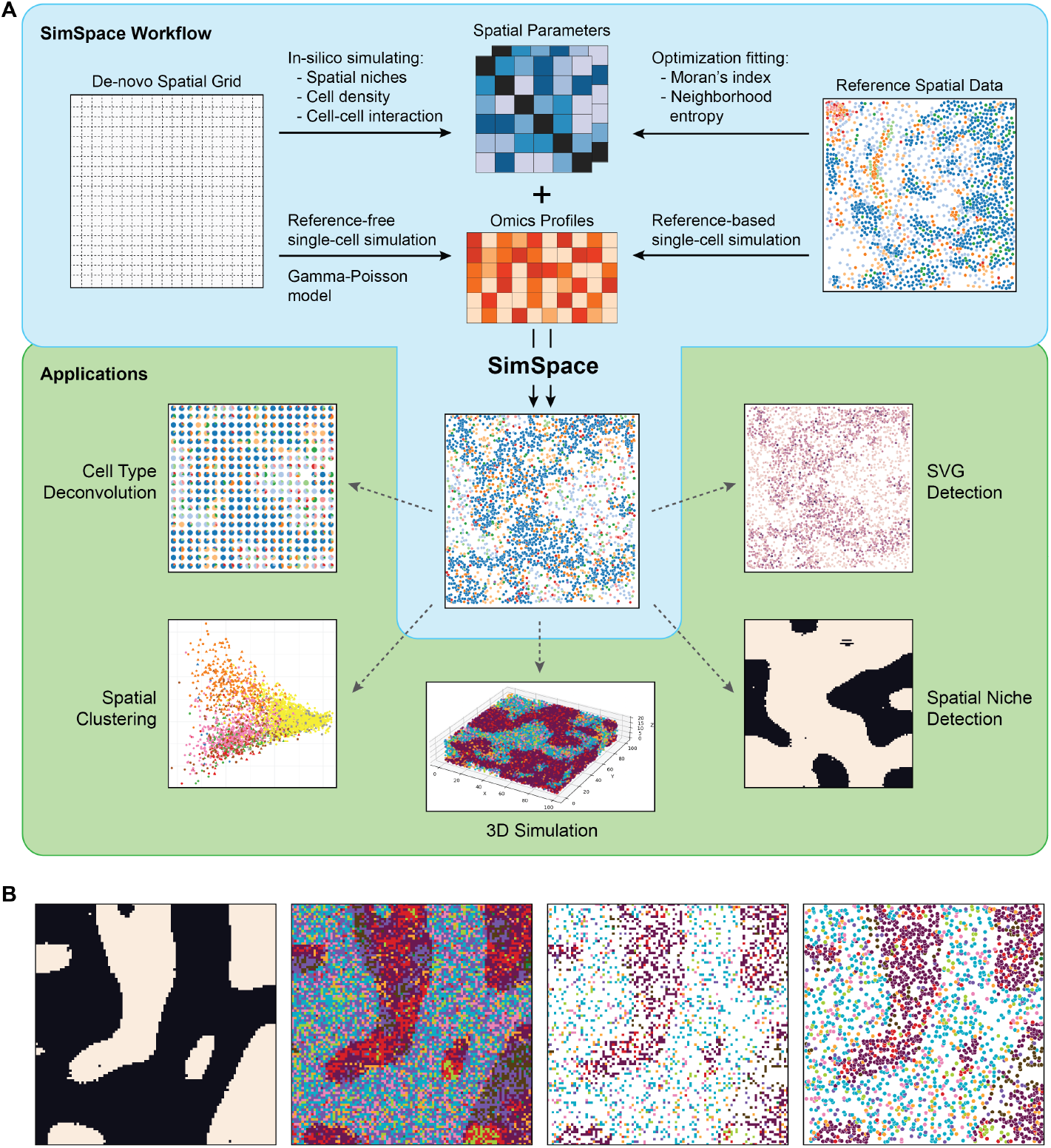
Overview of SimSpace framework with downstream applications and stepwise demonstration. (A) The SimSpace framework consists of two main components: the spatial simulator and the omics profile simulator. It supports both reference-free and reference-based simulations. For the spatial simulator, the framework first generates spatial grids with simulation parameters, controlling spatial domains, local cell-type organizations and densities, and cell-cell interaction patterns. These parameters are optionally fit to reference spatial datasets by optimizing spatial metrics. The molecular components are generated from built-in tools or external single-cell simulators, which will be integrated with spatial part as the final output. The resulting datasets enable various downstream applications, including spatial clustering, cell type deconvolution, SVG detection, spatial niche identification, and 3D spatial simulations. (B) A SimSpace simulation example workflow. First, the spatial domains are simulated using a niche-level MRF model, which generates spatial domains with different spatial structures and affinity patterns. Then, the cell types are simulated within each domain using a cell-level simulation model, which captures local cell-type organization and cell-cell interaction preferences. Then, cell densities are simulated, and a perturbation is applied to the cell coordinates to generate a continuous spatial distribution eventually.

SimSpace supports both reference-free and reference-based simulation. In the reference-free mode, users can specify tissue shape, spatial domain boundaries, niche affinities, cell-type affinities, phenotype-specific densities, marker-feature programs, and ligand–receptor-like spatial interaction effects. This enables controlled generation of diverse tissue architectures, including segregated domains, intermixed niches, spatial gradients, and localized molecular interaction patterns. In the reference-based mode, SimSpace calibrates its spatial parameters to an annotated spatial omics dataset by matching summary statistics such as spatial autocorrelation, neighborhood diversity, and cell-type composition. This likelihood-free calibration strategy preserves the interpretability of the generative model while allowing synthetic tissues to reproduce key spatial characteristics of real samples. SimSpace also supports three-dimensional simulation, enabling volumetric tissue layouts with controllable spatial organization.

We systematically evaluate SimSpace using complementary spatial statistics and benchmark its outputs against real datasets, including a Xenium human breast tumor dataset [33], a MERFISH mouse hypothalamus dataset [34], and a CODEX, also known as PhenoCycler-Fusion [35], human endometrial tumor dataset. These analyses show that SimSpace can reproduce multiple evaluated spatial characteristics, ranging from local spatial autocorrelation and neighborhood diversity to domain-level structure and gene-level distributional properties, across both reference-free and reference-guided settings. We further demonstrate its utility for benchmarking cell-type deconvolution, spatially variable gene detection, and in silico perturbation of cell–cell interactions. SimSpace is implemented as an open-source Python package with flexible parameterization and a modular design, enabling users to generate ground-truth-controlled synthetic tissues tailored to different biological and analytical questions. Together, these features make SimSpace a versatile resource for benchmarking, stress-testing, and developing computational methods for spatial omics.

## Results

### Overview of SimSpace

SimSpace is a flexible and comprehensive framework designed to simulate spatial omics data through a hierarchical generative strategy (**Supp Fig. 1**). The framework first generates a spatial tissue scaffold, which may be specified as a regular grid, an irregular mask, or a user-defined spatial domain. It then simulates tissue-scale spatial niches using a MRF, allowing users to control domain structure, and niche-level affinity patterns. Conditional on these niche labels, SimSpace assigns cell phenotypes through a second, niche-conditioned MRF that models spatial autocorrelation, local cell-type composition, and cell–cell interaction preferences within and across neighboring niches. Phenotype-dependent density parameters and coordinate jittering are then applied to obtain the final continuous spatial coordinates of individual cells (**Fig. 1B**).

For the molecular component, SimSpace generates cellular omics profiles conditional on the simulated cell phenotypes and spatial context. This can be achieved using built-in probabilistic models, including marker-feature programs, spatially variable features, and ligand–receptor-like spatial interaction modules, or by calling external single-cell simulation tools. The final output is a synthetic spatial omics dataset containing spatial coordinates, cell-type labels, omics profiles, and known ground truth spatial signals. Because the underlying tissue architecture, cell-type composition, spatial interaction structure, marker features, and spatially variable features are known by construction, SimSpace provides controlled ground truth for downstream applications such as cell-type deconvolution, spatially variable gene (SVG) detection, spatial clustering, and in silico perturbation analysis.

SimSpace’s spatial simulator is built on MRF models, a class of probabilistic graphical models widely used to represent local dependence among spatially indexed random variables [36–38]. In spatial omics, this local-dependence assumption is biologically natural: the phenotype composition and molecular state of a tissue region are often shaped more strongly by nearby cells than by distant cells (**Fig. 2A, Supp Fig. 2A**). By defining neighborhoods through a spatial graph, SimSpace can encode local proximity structure, niche organization, and cell-type co-localization in a statistically explicit and interpretable way.

**Figure 2.**
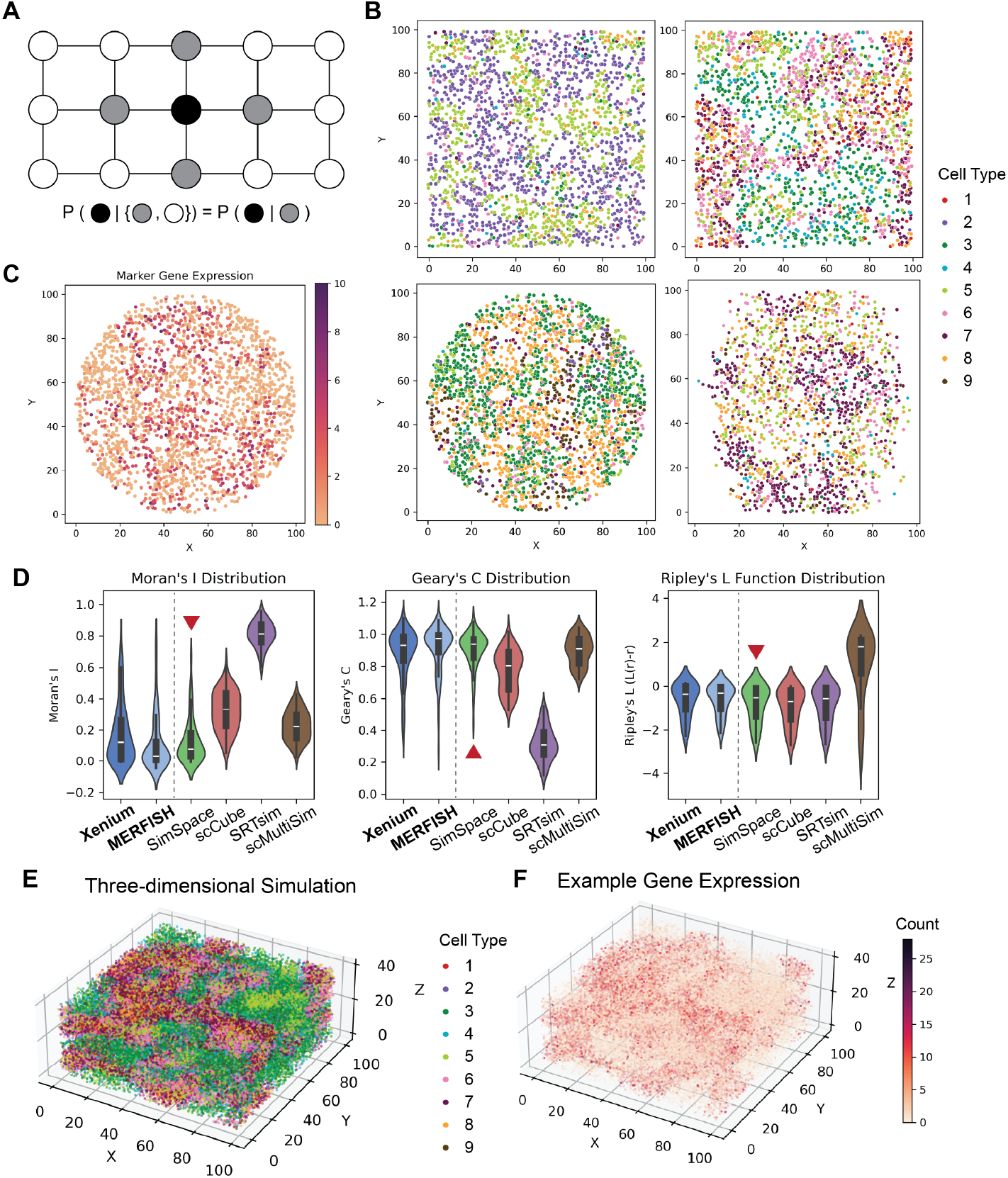
SimSpace enables the simulation of biologically realistic spatial patterns and molecular profiles in a reference-free setting. (A) Schematic illustrating the use of a MRF model in SimSpace to generate spatial patterns by enforcing local neighborhood dependencies, where the probability of a cell phenotype assignment is conditioned on its neighbors. (B) Representative examples of simulated two-dimensional spatial maps generated under varying spatial parameters, demonstrating the framework’s ability to produce diverse and tunable cell type arrangements; distinct cell types are indicated by different colors. (C) Simulated gene expression values for a representative marker gene display spatial gradients and local variability consistent with biological observations. (D) Comparative analysis of spatial patterns in real and simulated data using Moran’s Index score, Greary’s Contiguity ratio, and Ripley’s L function; SimSpace simulations (highlighted with red marks) more closely recapitulate the spatial patterns observed in the Xenium and MERFISH reference dataset than other reference-free simulation methods, including scCube, SRTsim, and scMultiSim. (E) Example of a three-dimensional spatial simulation generated by SimSpace, showing spatial niches across Z-slices annotated by cell type. (F) Simulated gene expression in three-dimensional space reveals spatial heterogeneity that mirrors the complexity of real tissue architecture.

A key feature of SimSpace is its two-level affinity structure. At the tissue scale, niche-affinity parameters control how spatial domains co-occur and segregate. At the cellular scale, niche-conditioned phenotype-affinity parameters control which cell types preferentially co-localize within or across neighboring niches. Together with modular omics generation components, this hierarchical design enables SimSpace to generate spatial architectures that are both controllable and biologically interpretable, rather than treating spatial coordinates, cell labels, and molecular profiles as largely separate simulation components.

SimSpace supports both reference-free and reference-based simulation. In the reference-free mode, users specify MRF parameters, cell densities, tissue-domain boundaries, and molecular simulation modules to generate de novo spatial omics datasets under controlled generative assumptions. In the reference-based mode, when an annotated spatial omics dataset is available, SimSpace calibrates the spatial parameters to match selected spatial features of the reference, such as cell-type composition, spatial autocorrelation, and neighborhood diversity, while still generating new spatial realizations that are not restricted to the original coordinate scaffold. This versatility enables SimSpace to produce a broad spectrum of synthetic spatial omics datasets across tissue geometries, resolutions, spatial organizations, and molecular modalities.

Despite its flexible modeling components, SimSpace remains computationally efficient. In runtime benchmarks (**Supp Fig. 1B**), SimSpace scaled approximately linearly with dataset size and maintained a low memory footprint (< 1 GB for up to 40,000 cells), comparable to or faster than existing reference-free simulators.

### Reference-free simulation of spatial omics data

SimSpace enables reference-free simulation of spatial omics data, allowing users to generate synthetic datasets with customizable spatial patterns and cell type distributions independent of any reference dataset. This mode is particularly advantageous when reference data are unavailable or when there is a need to model specific spatial architectures not observed in existing datasets. The reference-free simulation workflow begins with the specification of key hyperparameters, including the number of spatial niches, cell types, the neighborhood structure, and the spatial grid configuration. Using these parameters, SimSpace generates spatial cell maps exhibiting diverse and biologically relevant spatial organizations (**Fig. 2B**). Subsequently, omics profiles for individual cells are simulated in a reference-free manner (**Fig. 2C**), utilizing either the built-in Gamma-Poisson probabilistic model or external single-cell simulation tools such as splatter [39]. To incorporate spatial context into the simulated profiles, SimSpace offers an optional adjustment step based on a spatial ligand-receptor interaction model, which modifies the expression levels according to cell positions and phenotypes. Such interactions are known to play critical roles in spatial transcriptomics [40, 41]. This step enhances the biological realism of the reference-free simulated data by coupling molecular variation to spatial context. In addition to local interactions defined by the MRF, SimSpace allows users to impose larger-scale spatial structure by specifying spatial domains and the spatial interactions. User-defined spatial trends (e.g., along a proximal-distal axis, **Supp Fig. 2B-C**) can be overlaid on the MRF-generated cell map to introduce tissue-level heterogeneity that is not captured by local interactions alone. Collectively, the reference-free simulation mode provides a flexible platform for generating spatial omics datasets with user-defined spatial and phenotypic characteristics, facilitating method development and benchmarking in the absence of empirical references.

To assess the ability of SimSpace to generate realistic spatial organization in a reference-free setting, we generated eight simulated datasets using randomly sampled spatial parameters and compared them with other reference-free simulators and two real spatial omics datasets, a Xenium human breast tumor sample [33] and a MERFISH mouse brain sample [34]. We evaluated spatial structure using three widely used metrics including Moran’s I [42], Geary’s C [43], and Ripley’s L [44] function, which together capture spatial autocorrelation, local heterogeneity, and point-pattern clustering across cell types. For each cell type in the real and simulated datasets, we computed the distribution of each metric and compared these distributions across methods (**Fig. 2D**). Additionally, we also considered the cell-type interactions, which were measured by the cell-type interaction score from scFeatures [45,46] (**Supp Fig. 2D**). SimSpace produced distributions that closely resembled those of the Xenium and MERFISH references across all these metrics, indicating that the simulated tissues recapitulate key aspects of real spatial organization. In contrast, simulations generated using other reference-free spatial simulators, including scCube, SRTsim, and scMultiSim, showed substantial deviations from the distributions of real datasets, often displaying inflated autocorrelation or altered clustering behavior. These results indicate that, under the evaluated metrics and datasets, SimSpace more closely matches the reference distributions than the compared reference-free simulators.

### Three-dimensional spatial omics data simulation

SimSpace extends its simulation capabilities to three-dimensional spatial omics data, enabling the modeling of tissue architectures with increased biological realism. In this mode, the simulation framework generalizes the MRF model to three spatial dimensions, allowing users to specify a 3D spatial grid, including the number of layers and the neighborhood structure in all three axes. Both reference-free and reference-based simulation modes are supported, with spatial patterns, cell type distributions, and omics profiles generated in a manner analogous to the two-dimensional case. The resulting 3D spatial omics datasets capture complex spatial relationships and heterogeneity inherent to volumetric tissues (**Fig. 2E,F**). This functionality facilitates the benchmarking and development of computational methods designed for three-dimensional spatial omics data analysis.

### Reference-based simulation of spatial omics data

SimSpace further supports reference-based simulation of spatial omics data, in which synthetic datasets are generated to match selected spatial properties of a given reference sample while allowing new spatial realizations (**Fig. 3A**). In this mode, SimSpace calibrates the spatial parameters of the simulator by minimizing a loss function that compares spatial summary statistics between simulated and reference datasets. In the current implementation, the calibration loss includes Moran’s *I*, which measures cell-type spatial autocorrelation, and neighborhood entropy, which summarizes local cell-type diversity. The optimization is performed using a genetic algorithm that updates the niche-level affinity matrix, the niche-conditioned cell-type affinity parameters, phenotype-specific density parameters, and other user-specified spatial parameters. This likelihood-free calibration strategy allows SimSpace to reproduce selected spatial characteristics of the reference without requiring evaluation of the MRF normalizing constants. After calibration, SimSpace generates new spatial cell maps by running the calibrated niche-level and cell-phenotype MRFs, followed by the density and coordinate-jittering steps described above. When omics profiles are available in the reference dataset, SimSpace generates molecular measurements conditional on the simulated cell-type labels using built-in probabilistic models or external single-cell simulation tools such as scDesign3 [29]. The resulting dataset contains newly simulated spatial coordinates, cell-type annotations, and omics profiles, thereby providing a reference-guided but non-identical realization of the original tissue.

**Figure 3.**
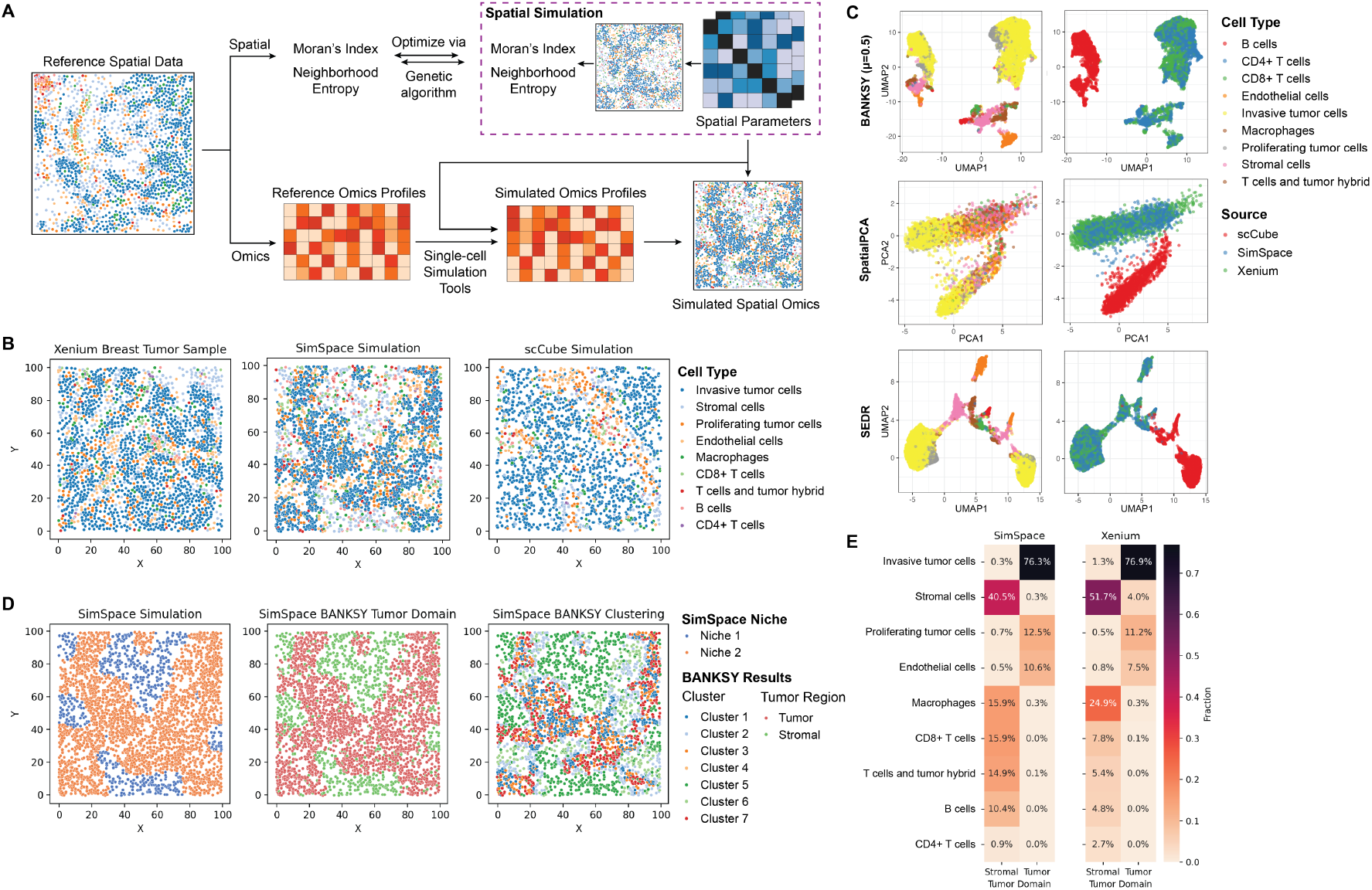
SimSpace enables reference-based spatial simulation and generates realistic, unbiased spatial omics data. (A) Reference-based simulation workflow. Given a reference spatial dataset, SimSpace uses a genetic algorithm to estimate the spatial parameters by matching spatial summary statistics between the simulated and reference configurations. These fitted spatial features are then combined with simulated omics profiles to generate realistic spatial omics datasets. (B) Cell type maps from real and simulated data. Reference tile from the Xenium breast tumor sample (left) is compared with simulations from SimSpace (middle) and scCube (right). SimSpace recapitulates key spatial organization features of the reference while introducing variability, unlike scCube which shows spatial patterns inconsistent with the reference. Evaluation using spatial clustering methods. Three tools (BANKSY, SpatialPCA, and SEDR) are applied to data from SimSpace, Xenium, and scCube. The left row shows clustering by cell type; the right row indicates data source. SimSpace simulations yield spatial embeddings that resemble those of the Xenium reference, yet do not cluster exclusively by source, suggesting that SimSpace simulations are both realistic and free of bias to the reference dataset. (D) Comparison of tumor-stromal domain structure in SimSpace simulations. The true niche structure encoded during simulation (left) is compared with BANKSY-derived spatial clusters (middle, right). BANKSY correctly identifies the major tumor and stromal domains and recovers the underlying spatial niche boundaries present in the SimSpace simulation. (E) Cell type composition across tumor and stromal domains. Proportions of each cell type in tumor versus stromal regions are shown for SimSpace simulations and the Xenium breast tumor reference. SimSpace accurately preserves key compositional differences between tumor and stromal compartments, including enrichment of invasive tumor cells in tumor regions and stromal and immune populations in stromal regions.

To evaluate the performance of SimSpace in reference-based simulation, we selected a 1mm × 1mm tile from the Xenium human breast tumor dataset [33] as the reference and generated a corresponding SimSpace dataset (**Fig. 3B**). While several existing tools, including scDesign3, SRTsim, and scCube, support reference-based simulation of gene expression, their spatial components typically reuse the original coordinates and assign simulated profiles back to these reference-derived coordinate scaffold. As a result, although these tools can generate realistic molecular profiles, they do not directly address the task of generating new spatial arrangements that are calibrated to the spatial properties of a reference tissue. In contrast, SimSpace uniquely performs reference-guided de novo spatial simulation by calibrating its generative spatial model to reference summary statistics, enabling new spatial configurations that are not tied to the reference geometry. Because available simulators do not solve exactly the same reference-guided spatial generation task, there is no direct apples-to-apples baseline for this setting. As a best-effort comparison, we report results from scCube using its reference-based expression simulation together with its reference-free spatial sampling procedure. This comparison illustrates the gap between reference-free coordinate sampling and reference-guided spatial modeling when doing reference-based simulations.

We evaluated whether the simulated datasets recapitulated key spatial and molecular characteristics of the reference sample. We applied spatial clustering methods, including BANKSY [21], SpatialPCA [22], and SEDR [47], to the reference and simulated datasets. The results showed that SimSpace-generated data aligned closely with the reference in these analyses, with reduced dataset-specific separation relative to the comparator simulation (**Fig. 3C, Supp Fig. 4A**). Particularly, scCube simulations exhibited a clear batch effect. To distinguish the contribution of spatial organization from that of omics-profile simulation, we performed a matched-profile control in which the simulated omics profiles from SimSpace and scCube were replaced by Xenium reference profiles sampled from cells with the same phenotype, while retaining the simulated coordinates and cell-type labels. The clustering patterns remained consistent under this control, indicating that the observed differences, especially the substantial batch effect in scCube’s data, were primarily driven by spatial organization rather than by molecular-profile simulation alone (**Supp Fig. 4B**). SimSpace also preserves gene-level distributional properties of the reference dataset, including the mean– variance relationship and the fraction of zeros per gene (**Supp Fig. 4D**).Because cell-phenotype clustering differs from detecting higher-order spatial domains, we further assessed whether SimSpace preserves tumor– stromal architecture. Using BANKSY, we found that the inferred tumor and stromal regions closely matched the niches encoded during simulation (ARI = 0.69, NMI = 0.59; **Fig. 3D, Supp Fig. 4C**), and the domain-specific cell-type compositions were strongly correlated with those of the Xenium reference (tumor: *r* = 0.99; stromal: *r* = 0.92; **Fig. 3E**). Together, these results show that SimSpace can reproduce the evaluated spatial organization and gene-level distributional characteristics of a reference tissue while generating new spatial realizations. Since the available comparators do not solve the same reference-guided de novo spatial generation task, these results should be interpreted as evidence of practical utility rather than a definitive same-task ranking across simulators.

**Figure 4.**
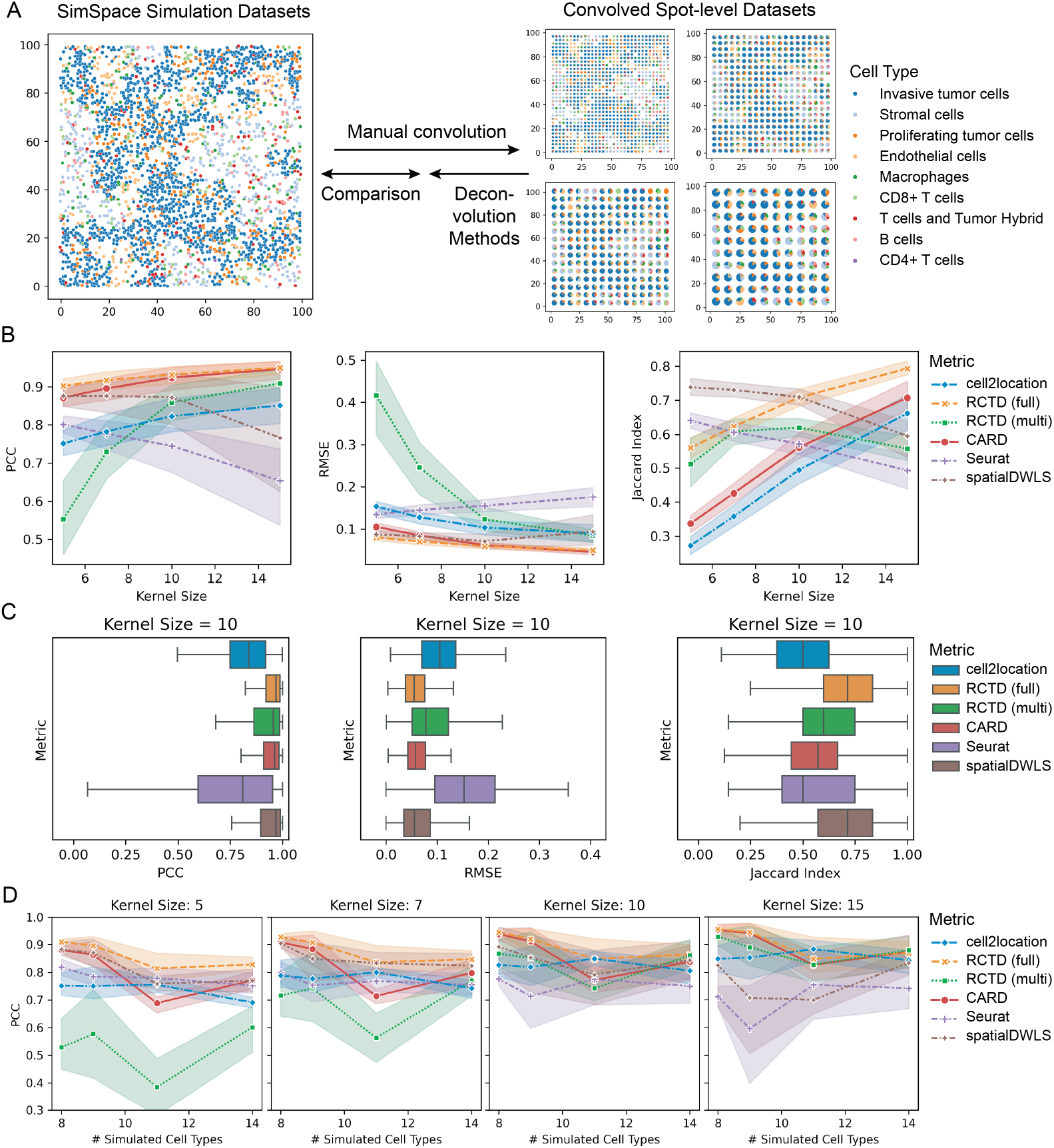
Benchmarking spatial cell type deconvolution methods using SimSpace simulations. (A) Bench-marking framework. Simulated single-cell resolution data are aggregated into spot-level measurements via manual convolution with varying kernel sizes to mimic real spatial transcriptomics. These convolved datasets serve as ground truth for evaluating deconvolution methods. (B) Performance across deconvolution methods evaluated using three metrics, PCC, RMSE, and Jaccard index, across kernel sizes (spot resolutions). SimSpace simulations reveal performance trends across methods, including strong performance from RCTD and CARD. (C) Comparison at fixed kernel size (10). Boxplots summarize PCC, RMSE, and Jaccard index across all methods. CARD and cell2location achieve high accuracy and consistency across metrics. (D) Robustness across varying cell type complexity. Performance of each method is shown as a function of the number of simulated cell types under four different kernel sizes. RCTD full mode, CARD, and cell2location demonstrate robustness across increasing complexity, while performance for other methods varies more widely.

### Benchmarking computational methods using SimSpace simulated datasets

SimSpace generates spatial omics datasets with user-controllable spatial architecture, cell-type composition, molecular signals, and known ground truth. This makes it useful as an in silico platform for spatial omics computational methods under controlled and reproducible settings. Here, we illustrate this utility for two common spatial omics tasks: cell-type deconvolution and spatially variable gene detection.

#### Benchmark interpretation

The benchmarking exercises in this study are intended to evaluate whether SimSpace-generated datasets can support controlled assessment of downstream methods. They should not be interpreted as exhaustive rankings of all available analysis tools, as the performance depends on dataset design, reference-tile selection, spatial resolution, preprocessing choices, and each method’s modeling assumptions. Similarly, for reference-guided simulator comparisons, existing tools do not all solve the same task as SimSpace; therefore, those comparisons should be viewed as practical context rather than as a strict leaderboard.

#### Cell type deconvolution

Cell type deconvolution is a fundamental task in spatial transcriptomics analysis, aiming to estimate the relative abundance of distinct cell types within each spatial location, particularly for technologies with multicellular resolution [48, 49]. We evaluated five widely used cell type deconvolution methods: cell2location [7], RCTD (in both ‘full’ and ‘multi’ modes) [8], CARD [9], Seurat [10], and spatialDWLS [11]. These methods represent state-of-the-art approaches for estimating cell-type composition in spatial omics data and have been extensively benchmarked in recent studies [50, 51].

To benchmark these methods under controlled ground truth, we used SimSpace to generate reference-based simulated datasets from randomly selected tiles of the Xenium human breast tumor dataset. To span a broad range of biological and spatial complexity, we varied the number of cell types and the spatial patterns for each reference tile, resulting in 16 distinct simulated datasets (**Supp Fig. 5**). Each dataset was further processed using four different convolution kernel sizes, thereby mimicking different spatial resolutions. Deconvolution methods were then applied to these convolved datasets, and predicted cell-type proportions were compared with the known proportions from the original SimSpace simulations (**Fig. 4A**). Evaluation metrics included Pearson correlation coefficient (PCC), root mean square error (RMSE), and Jaccard Index (JI), were calculated for each convolved spot to quantitatively assess the accuracy of cell type proportion estimates.

**Figure 5.**
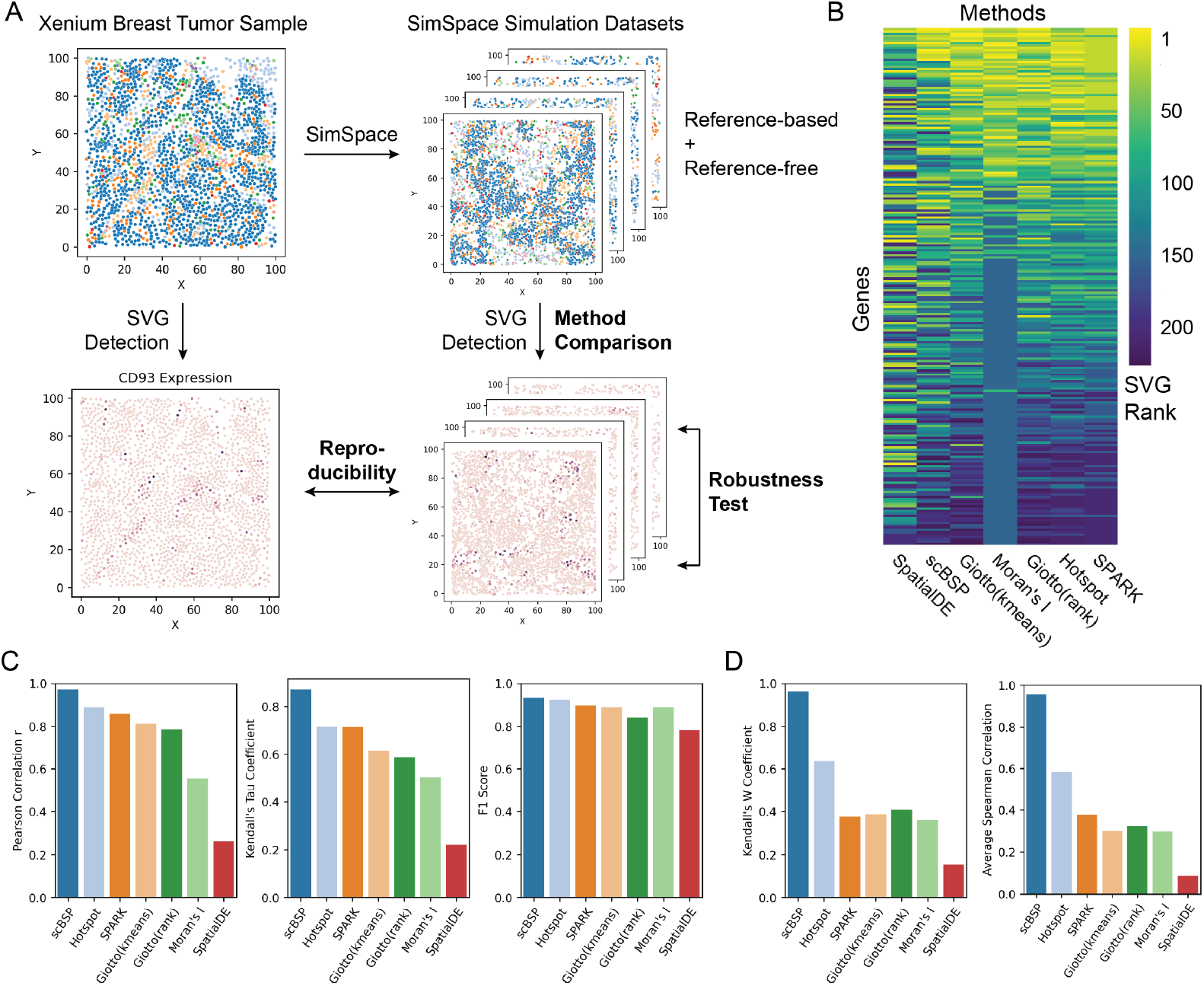
Benchmarking SVG detection using SimSpace simulations. (A) Benchmarking framework. A Xenium human breast tumor sample is used to generate SimSpace datasets under both reference-based and reference-free simulation modes. SVG detection is performed on the original Xenium dataset and on simulated datasets to compare methods and assess reproducibility and robustness. (B) Comparison of SVG rankings across methods. Rank-normalized p-values from a reference-based SimSpace simulation are visualized across seven methods: SPARK, Hotspot, Giotto (rank), Moran’s I, Giotto (k-means), scBSP, and SpatialDE. Although top-ranked genes show general overlap, several methods like scBSP and SpatialDE shows discrepancies in gene prioritization. (C) Reproducibility between simulated and reference data. Across methods, Pearson correlation (left), Kendall’s Tau (middle), and F1 score (right) are computed between the Xenium reference and reference-based SimSpace simulation. Most methods demonstrate high reproducibility, with scBSP achieving the strongest concordance across all metrics. (D) Robustness across reference-free simulations. Consistency of SVG rankings across eight reference-free SimSpace datasets is quantified using Kendall’s W (left) and average Spearman correlation (right). scBSP and Hotspot show the highest robustness to variations in spatial structure, while other methods display more variability.

The results indicate that most deconvolution methods achieve high PCC with the ground truth cell type proportions, with RCTD and CARD consistently showing the strongest overall performance (**Fig. 4B, C, Supp Fig. 6**). Similar trends were observed for RMSE. Notably, while most methods exhibit improved performance with increasing convolution kernel sizes, Seurat and spatialDWLS performed best at smaller kernel sizes, suggesting that these methods may be more sensitive to spatial resolution and potentially less suitable for datasets with larger spot sizes, such as Visium. To further assess the ability of each method to detect the presence of specific cell types, we evaluated performance using JI. Consistent with the PCC and RMSE results, Seurat and spatialDWLS achieve higher JI at smaller kernel sizes, while several other methods improved with larger kernels. We also varied the number of cell types from 8 to 14 and found that all methods tended to perform better when fewer cell types were present (**Fig. 4D**), consistent with the increased difficulty of deconvolving more heterogeneous mixtures. Small local performance fluctuations at specific cell-type numbers, such as 9 − 11 cell types, likely reflect stochastic variation in simulation, convolution, and sparse or imbalanced spot compositions, and did not alter the overall performance trends. In summary, our benchmarking of cell type deconvolution methods using SimSpace-simulated datasets demonstrates that most approaches are capable of accurately recovering cell type proportions across a range of spatial resolutions and complexities. Among the evaluated methods, RCTD and CARD consistently exhibited the highest accuracy across varying kernel sizes. Additionally, we observed that increasing the number of cell types in the dataset led to a reduction in deconvolution accuracy for all methods, reflecting the greater challenge posed by higher cellular complexity. These results underscore the value of SimSpace for generating controlled, ground-truth datasets that enable systematic and rigorous evaluation of spatial deconvolution algorithms under diverse experimental conditions.

**Figure 6.**
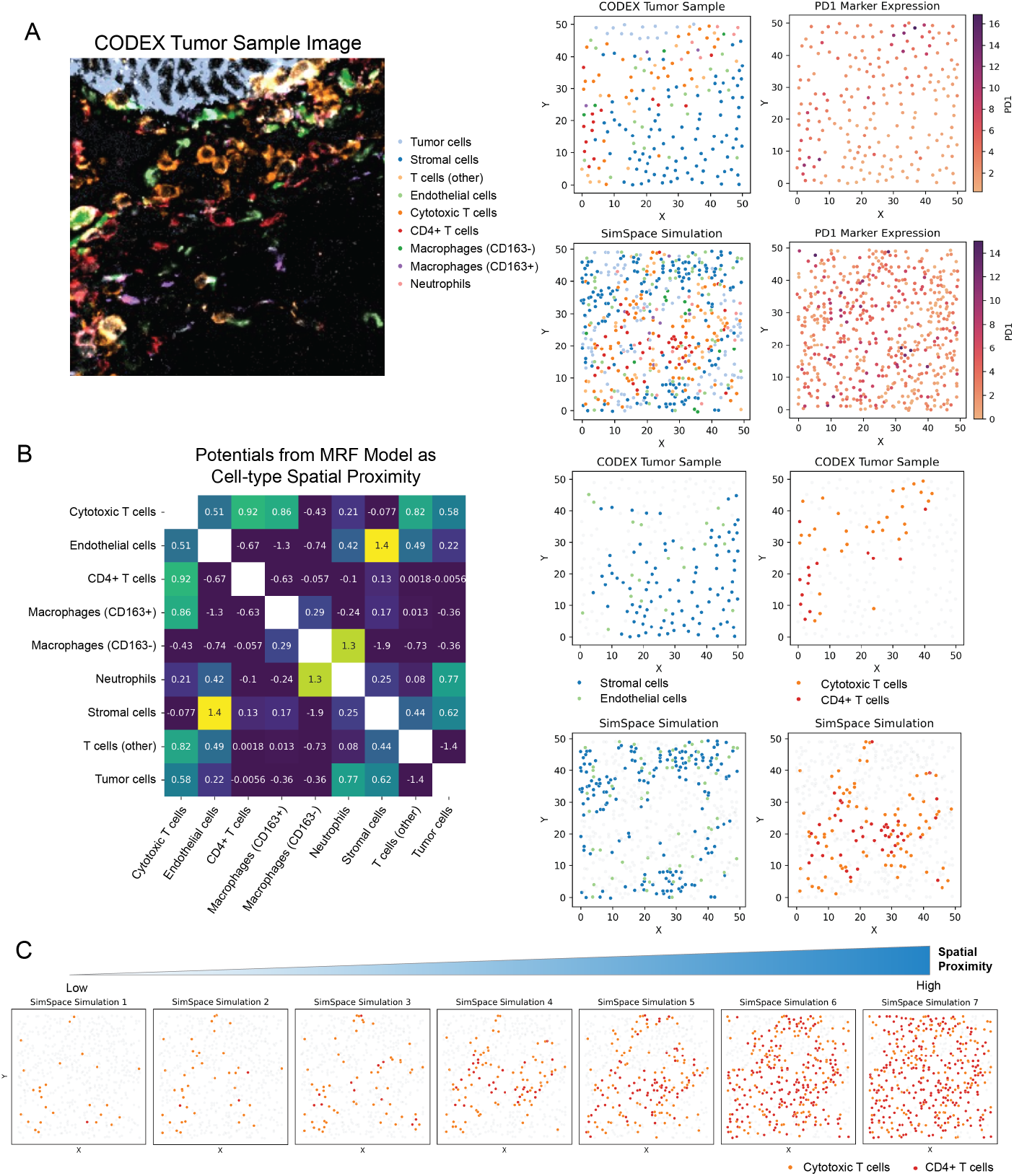
SimSpace models and simulates spatial proteomics data to reveal microenvironmental cell-cell interactions. (A) Reference-based simulation of spatial proteomics. A representative tile from a CODEX endometrial tumor sample (left) is used to generate a SimSpace simulation incorporating both spatial cell distributions and protein marker expression. PD1 expression is shown for both real and simulated datasets, demonstrating the preservation of spatial expression patterns across cell types. (B) Quantifying spatial cell-cell interactions. Using SimSpace’s underlying MRF model, an interaction matrix is inferred to represent pairwise spatial co-localization strengths between cell types. Strong spatial attraction is observed between cytotoxic and CD4+ T cells, and between endothelial and stromal cells, while tumor cells show spatial exclusion from most immune populations. Spatial distributions of selected cell types (endothelial and stromal cells, cytotoxic and CD4+ T cells) are shown in both the CODEX reference and SimSpace simulation, demonstrating realistic tissue architecture reconstruction. (C) In-silico perturbation experiments using SimSpace. By modifying the interaction parameters in the MRF model, SimSpace can simulate hypothetical scenarios of altered cell-cell interactions. Here, we show the effect of increasing attraction between CD4+ cells and cytotoxic T cells, resulting in enhanced spatial co-localization in the simulated tissue architecture. This capability allows researchers to explore potential therapeutic strategies and their impact on the tumor microenvironment in a controlled computational setting.

#### Spatially variable gene detection

SVG detection is another common task in spatial transcriptomics analysis, where the goal is to identify genes that exhibit non-random spatial structure across a tissue section [52, 53]. We compared six different SVG detection methods, including Hotspot [12], SpatialDE [13], scBSP [14, 15], SPARK [16], Giotto [17], and Moran’s I as implemented in Seurat [54].

We generated SimSpace datasets from the Xenium human breast tumor reference under both reference-based and reference-free settings (**Fig. 5A**). For the reference-based scenario, the simulated dataset was calibrated to reproduce selected spatial and gene-expression characteristics of the Xenium reference, allowing us to assess the reproducibility of SVG detection between real and reference-guided synthetic data. In parallel, in the reference-free setting, we generated eight additional datasets with varying spatial patterns while using reference-derived gene-expression profiles. SVG detection methods were applied to these datasets to assess the robustness of each approach and to investigate the impact of varying spatial patterns on SVG detection. Because SVG detection methods use different statistical models and return p-values with different scales and distributional properties, direct comparison of raw p-values can be misleading. We therefore used a rank-based normalization procedure: within each method and dataset, genes were ranked by statistical significance, with smaller p-values corresponding to stronger spatial signal, and ranks were mapped to a common scale [55]. This normalization enabled method outputs to be compared across datasets and across methods. Using the reference-based simulated dataset, we assessed the concordance of normalized p-values among the different methods (**Fig. 5B**). While there was general agreement in the identification of significant genes, notable discrepancies in gene rankings were observed, particularly for SpatialDE and scBSP, which exhibited rankings distinct from those produced by the other methods. These differences underscore the impact of methodological choices on SVG detection outcomes.

To further evaluate reproducibility, we next compared the rank-normalized p-values obtained from the reference-based simulated dataset to those from the original Xenium reference dataset. With the exception of SpatialDE and Moran’s Index, most methods demonstrated strong concordance between simulated and reference data, as reflected by PCC ranging from 0.6 to 0.9 (**Fig. 5C, Supp Fig. 7**). Kendall’s tau, a rank-based correlation measure, showed consistent trends. Notably, scBSP achieved the highest correlation with the reference dataset. To assess binary detection performance, we identified genes as significant SVGs using a *p*-value threshold of 0.05 and quantified overlap between simulated and reference datasets. Most methods achieved high precision and recall, with F1 scores between 0.8 and 0.9 (**Fig. 5C**). Collectively, these findings demonstrate that most SVG detection methods yield highly reproducible results between SimSpace-generated simulated data and real datasets, underscoring the utility of SimSpace as a robust simulation platform for benchmarking spatial analysis tools.

For the reference-free simulations, we applied the same rank-based normalization procedure to the p-values generated by each method and evaluated the stability of SVG detection across independently generated datasets (**Fig. 5D**). We quantified concordance using Kendall’s coefficient of concordance *W* and average Spearman correlation. Among the methods evaluated, scBSP exhibited the highest level of consistency across reference-free simulations, indicating robustness to variations in spatial patterns. Hotspot also demonstrated substantial concordance, whereas the remaining methods showed greater sensitivity to changes in spatial organization. Thus, our benchmarking of SVG detection methods using SimSpace-simulated datasets demonstrates that most approaches yield highly reproducible and robust results when compared to real spatial omics data. Notably, methods such as scBSP and Hotspot exhibited the highest levels of consistency and robustness to variations in spatial patterns, These results highlight the utility of SimSpace for generating controlled, ground-truth spatial omics datasets, thereby enabling systematic evaluation and comparison of SVG detection algorithms and supporting the advancement of spatial analysis methodologies.

In summary, the SVG benchmarking analyses show that SimSpace can support both reference-guided reproducibility assessment and reference-free stress-testing of spatial analysis methods. Importantly, these analyses should be interpreted as evaluating method behavior under the simulated scenarios considered here, rather than as a universal ranking of SVG detection tools.

### SimSpace for spatial proteomics data simulation

Beyond spatial transcriptomics, SimSpace can also be applied to simulate spatial proteomics data, where molecular measurements consist of protein-marker intensities rather than transcript counts. This capability is important for studying the spatial organization of tissue microenvironments, especially in imaging-based assays that jointly measure cellular phenotype, protein state, and spatial context [56]. To demonstrate this, we applied SimSpace to a selected tile of a CODEX (CO-Detection by indEXing [57]) human endometrial tumor dataset, which contains spatial proteomics data with 31 protein markers and 206 cells with 9 phenotypes. We generated a reference-based simulated dataset based on the reference tile from the CODEX dataset, and then applied SimSpace to simulate the spatial patterns of the tumor samples and the corresponding protein marker expression profiles in the tumor microenvironment (**Fig. 6A**).

By leveraging the MRF model, SimSpace encodes the spatial organization of the tissue into affinity parameters, which summarize conditional spatial proximity preferences between cell types (**Fig. 6B**). Positive affinity values indicate that two cell types are more likely to occur as neighbors under the fitted spatial model, whereas negative values indicate reduced local adjacency. These parameters should be interpreted as model-based interaction affinities rather than direct measurements of physical or causal cell–cell interactions. In the CODEX example, the calibrated affinity matrix suggested strong local co-occurrence between cytotoxic T cells and CD4^+^ T cells (*θ* = 0.92), as well as between endothelial and stromal cells (*θ* = 1.40) (**Fig. 6B**), suggesting that these cell types tend to co-localize in the microenvironment. The SimSpace-generated tissue recapitulated these qualitative spatial patterns, including clustering of T-cell subsets and proximity between endothelial and stromal cells, consistent with known tumor-microenvironment organization reported in related studies [58, 59]. This underscores SimSpace’s capacity to generate biologically realistic spatial architectures based on inferred affinity parameters. Conversely, tumor cells showed negative affinities with most immune cell types, except neutrophils and cytotoxic T cells, suggesting a distinct spatial organization consistent with reported features of endometrial tumor microenvironments [60, 61].

To illustrate how the fitted affinity parameters can be used for interpretable in silico perturbation, we systematically increased the interaction strength between cytotoxic T cells and CD4^+^ T cells while keeping other simulation components fixed. Stronger positive affinity produced progressively tighter co-localization of these T-cell subsets (**Fig. 6C**). This experiment demonstrates how SimSpace can generate model-based counterfactual tissue architectures by modifying specific entries of the interaction matrix. Such perturbations provide a controlled way to explore how changes in local cellular affinities, such as enhanced immune co-localization or infiltration, could reshape the spatial structure of the tumor microenvironment. Together, these analyses support SimSpace as an extensible framework for spatial proteomics simulation and for interpretable perturbation experiments based on modifiable spatial interaction parameters.

## Discussion

Our intended use case for SimSpace is methodological evaluation rather than substitution for biological validation. In particular, realism in this study is operationalized through the spatial statistics, domain-level structure, and downstream benchmarking behaviors that we explicitly evaluate. The framework is therefore strongest as a resource for controlled benchmarking, stress-testing, and counterfactual simulation, while questions requiring full tissue fidelity must still be addressed in real data. Simulating spatial omics data remains an essential but unresolved challenge. While simulated data cannot replace real tissues for interrogating biological relevance, they serve a critical complementary role by providing ground-truth-controlled settings for evaluating method behavior. In this study, we introduce SimSpace, a unified framework that addresses several limitations of existing simulation approaches by generating spatially structured tissues and molecular profiles in both reference-free and reference-guided modes.

A central contribution of SimSpace lies in its reference-based spatial modeling strategy. Existing reference-based simulators, including scDesign3, SRTsim, and scCube, reuse the spatial coordinates of the input tissue and add variation only at the omics level. As a result, they do not learn or regenerate the spatial dependencies, niche boundaries, or interaction patterns inherent in the reference. In contrast, SimSpace learns spatial priors from the reference dataset through its MRF formulation—capturing spatial domains, spatial autocorrelation, and pairwise cell-cell spatial proximity—and then generates entirely new tissue architectures that preserve the underlying biological principles without replicating the original geometry. This enables SimSpace to produce novel spatial configurations informed by the reference data, supporting benchmarking and in-silico perturbation studies.

Across the datasets and spatial metrics evaluated here, we show that SimSpace simulations can match several key measured spatial properties of the reference tissues. Expanded benchmarking demonstrates that SimSpace reproduces global and local spatial dependencies more accurately than other reference-free simulators, including scCube, SRTsim, and scMultiSim. In the reference-based setting, SimSpace additionally preserves higher-order spatial domains, such as tumor-stromal architecture, with strong agreement in both spatial boundaries and domain-specific cell-type compositions. At the molecular level, SimSpace captures gene-level distributional properties, including mean-variance structure and dropout patterns, consistent with the Xenium reference data. Together, these results support SimSpace as a practical framework for modeling spatial organization and molecular variation for benchmarking applications.

The hierarchical structure of SimSpace is another major strength. By decoupling (i) niche structure, (ii) cell-type spatial interactions, and (iii) omics profile generation, SimSpace allows users to independently tune spatial heterogeneity, cellular proximity, density, and expression distributions. This modularity also enables in-silico perturbation experiments by systematically modifying *θ* to explore hypothetical changes in cell-cell interactions, as demonstrated in our spatial proteomics analysis of CODEX endometrial tumors. Because SimSpace learns interaction strengths directly from reference data, these perturbations are interpretable and biologically grounded.

Despite these capabilities, several limitations remain. First, because MRFs model only local spatial dependencies, they cannot inherently generate global structures such as proximal-distal gradients or long-range tissue axes. SimSpace partially addresses this through user-defined spatial domains and optional global expression trends, but modeling continuous within-cell-type spatial gradients remains an opportunity for future development. Second, although reference-based simulations incorporate spatial covariates into the omics generative model (via scDesign3), reference-free omics do not automatically encode spatial dependencies unless specified through ligand-receptor modules or domain structures. Finally, the current version of SimSpace operates at the single-sample level. Extending the framework to simulate multi-sample or multi-condition datasets (with controlled inter-sample variability, batch structure, or condition-specific spatial changes), as well as the multi-layer three-dimensional spatial datasets, would significantly expand its utility, particularly for benchmarking integration and differential spatial analysis methods.

In summary, SimSpace provides a flexible platform for spatial omics data simulation. By generating reproducible, ground-truth-controlled datasets that capture user-specified and reference-guided spatial structure, SimSpace enables benchmarking of computational tools and supports exploratory modeling of spatial biology. As spatial omics technologies continue to evolve, frameworks like SimSpace may play an increasingly important role in method development, hypothesis testing, and reproducible evaluation.

## Method

### SimSpace framework

The SimSpace model simulates spatial omics data through a hierarchical generative framework. Let *σ* denote a cell’s omics profile, *c* its cell phenotype or cell type, and (*x, y*) its spatial coordinate. We factorize the joint distribution of (*σ, c*, (*x, y*)) as

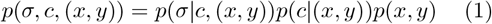

where *p*(·) denotes a generic probability distribution. This factorization corresponds to a natural simulation hierarchy. First, cell locations are generated from the spatial distribution *p*(*x, y*), which may be specified through user-defined spatial patterns and cell-density parameters. Second, cell phenotypes are generated from *p*(*c* | (*x, y*)), which allows the cell-type composition to vary across space and can be modeled, for example, using a MRF to capture spatial autocorrelation and interactions among neighboring cell types. Finally, omics profiles are generated from *p*(*σ* | *c*, (*x, y*)), allowing molecular measurements to depend on both the cell phenotype and spatial context. This final conditional distribution may be specified using reference-based single-cell omics simulators or reference-free probabilistic models.

Each component in this hierarchy can be specified flexibly to encode distinct biological assumptions, tissue architectures, or experimental designs. Following the factorization in (1), SimSpace generates synthetic spatial omics data by first sampling cell locations, then assigning cell types conditional on these locations, and finally generating omics profiles conditional on both cell type and spatial context:

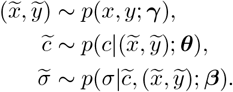

Here, 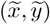 denotes the simulated spatial coordinate, 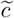 the simulated cell type or phenotype, and 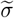 the simulated omics profile. The parameter vectors ***γ, θ***, and ***β*** govern the spatial distribution, spatially varying cell-type composition, and conditional omics-profile distribution, respectively.

### Spatial modeling of cell phenotypes using MRFs

To generate spatially structured cell phenotypes and tissue niches, SimSpace models the phenotype configuration, i.e., 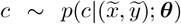, using a MRF. This choice provides a flexible probabilistic mechanism to encode local spatial dependence, while allowing users to specify or learn biologically meaningful cell–cell affinity patterns. As a probabilistic graphical model that represents the dependencies between random variables, MRFs have been widely used to model spatially dependent discrete states in image analysis, computer vision, and spatial omics settings [37, 62–64].

Let ℒ = {1, …, *N*} denote the set of simulated cell locations, with cell *i* located at *s*_*i*_ := (*x*_*i*_, *y*_*i*_). Let *c*_*i*_ ∈ {1, …, *K*} denote the phenotype or cell type assigned to location *s*_*i*_, where *K* is the number of cell types. A phenotype configuration is denoted by **c** = (*c*_1_, …, *c*_*N*_). To encode local spatial dependence, we construct an undirected neighborhood graph

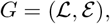

where (*i, j*) ∈ ℰ if cells *i* and *j* are spatial neighbors, for example if ∥*s*_*i*_ − *s*_*j*_∥ ≤ *r* for a user-specified distance threshold *r*, or if *j* belongs to the *K*_nn_-nearest-neighbor set of *i*. We write 𝒩_*i*_ ={ *j* : (*i, j*) ∈ ℰ} for the neighbors of cell *i*.

Under the MRF model, the phenotype at each location depends on the remaining phenotypes only through its spatial neighbors. That is,

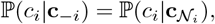

where **c**_−*i*_ denotes all phenotype labels except *c*_*i*_, and 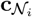 denotes the labels of the neighboring cells. This local Markov property [65] (**Fig. 2A**) is well suited to spatial omics data, where the phenotype composition of a local tissue region is often shaped more strongly by nearby cellular interactions than by distant cells. Studies have shown that such neighborhood definitions are effective in capturing local spatial relationships especially in spatial analysis [63, 64].

We specify a pairwise MRF for the joint phenotype configuration:

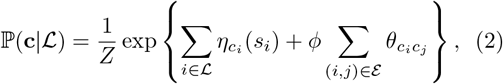

where *Z* is the normalizing constant,

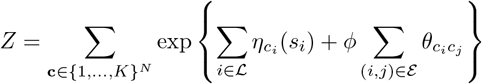

Here, *η*_*k*_(*s*_*i*_) is a cell-type-specific baseline term that allows the relative abundance of phenotype *k* to vary across space. The matrix **Θ** = (*θ*_*ab*_)_1≤*a,b*≤*K*_ encodes pairwise affinities between cell types: positive values favor local co-occurrence of phenotypes *a* and *b*, whereas negative values discourage such colocalization. The scalar *ϕ* ≥ 0 is a global smoothness parameter that controls the overall strength of spatial dependence, reflecting the degree of interaction between neighboring phenotypes. Larger values of *ϕ* generate smoother or more strongly segregated tissue domains, depending on the structure of Θ.

The conditional distribution needed for simulation follows directly from (2). For each cell *i* and candidate phenotype *k*,

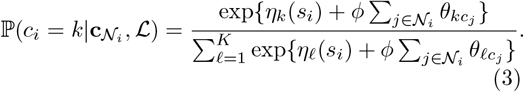

This formulation separates three biologically interpretable components: the spatial baseline abundance *η*_*k*_(*s*_*i*_), the cell-type affinity matrix **Θ**, and the global interaction strength *ϕ*.

The model parameters can either be learned from reference data or specified by the user based on prior biological knowledge. In reference-free settings, the baseline functions and affinity matrices may be randomly initialized or manually specified to generate controlled spatial patterns, such as clustered niches, segregated tissue domains, or enriched co-localization of selected cell-type pairs. SimSpace then uses Gibbs sampling to generate phenotype configurations from the specified MRF. Starting from an initial assignment, the sampler repeatedly updates each cell label conditional on the current labels of its neighbors. The procedure is summarized as follows:

1. Initialize phenotypes 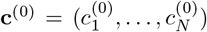, either at random or from a user-specified composition.
2. For each Gibbs sweep *t* = 1, …, *T*, visit cells *i* = 1, …, *N* sequentially. When cell *i* is updated, let 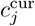 denote the current label of neighbor *j*, using the most recently available labels within the sweep. Compute

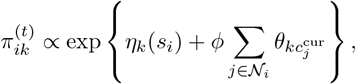

for *k* = 1, …, *K*, and normalize so that 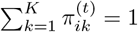.
3. Sample a new label for cell *i* from the categorical distribution with probabilities 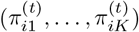.
4. After burn-in, return the final phenotype configuration, or retain multiple independent samples to quantify simulation variability.

This MRF-based simulator is flexible in both reference-based and reference-free settings. When annotated spatial or single-cell data are available, the baseline terms *η*_*k*_(*s*) and affinity parameters **Θ** can be estimated from the reference data. When such information is unavailable, they can be specified by the user to generate controlled spatial patterns, such as clustered niches, segregated tissue domains, or enriched co-localization of selected cell-type pairs. Thus, the MRF component provides an inter-. pretable and tunable mechanism for generating spatially structured cell-type compositions within the broader SimSpace generative framework.

### Reference-free simulation

SimSpace supports fully reference-free simulation, in which the user specifies the number of spatial niches, cell phenotypes, spatial domain, neighborhood structure, and omics-feature architecture without requiring an annotated reference dataset. This mode is designed to generate controlled spatial omics data with interpretable sources of variation, including tissue-domain organization, niche-specific cell-type composition, phenotype-dependent density, marker-feature expression, and spatially mediated ligand–receptor effects.

Let Ω ⊂ ℝ^2^ denote the spatial domain and let ℒ_0_ = {1, …, *N*_0_} denote an initial set of candidate lattice locations with coordinates 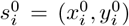. SimSpace allows the spatial domain to be a full rectangular grid, a binary mask, or a user-defined boundary function specifying the admissible tissue region. This flexibility enables the generation of irregular tissue outlines, partially occupied domains, layered structures, and spatial gradients (**Supp Fig. 2A**). On the admissible locations, we construct an undirected neighborhood graph

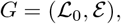

where (*i, j*) ∈ ℰ if locations *i* and *j* are spatial neighbors, for example according to a distance threshold or a *K*_nn_-nearest-neighbor rule. We write 𝒩_*i*_ ={ *j* : (*i, j*) ∈ ℰ for the neighbor set of location *i*. Larger neighborhoods correspond to longer-range local interactions and produce smoother spatial patterns.

#### Niche simulation

We first simulate a spatial niche label *h*_*i*_ ∈ {1, …, *M*} for each candidate location, where *M* is the number of tissue niches. Let 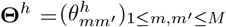 denote a symmetric niche-affinity matrix. The niche configuration 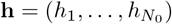 is generated from a pairwise MRF,

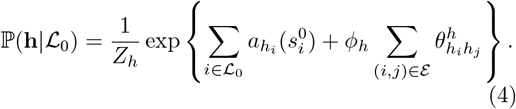

Here, *Z*_*h*_ is the normalizing constant, 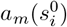 controls the spatial baseline prevalence of niche 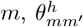 encodes the affinity between neighboring niches *m* and *m*′, and *ϕ*_*h*_ ≥ 0 controls the overall strength of niche-level spatial dependence. Positive affinity promotes local co-occurrence of the corresponding niche pair, whereas negative affinity discourages adjacency. In the absence of prior information, **Θ**^*h*^ and *a*_*m*_(·) can be specified by the user or generated randomly to create diverse tissue-domain patterns.

A Gibbs sampler is used to draw **h** from (4). Specifically, the conditional distribution for updating location *i* is

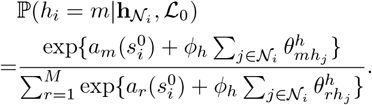

#### Cell-phenotype simulation

Conditional on the simulated niche labels, SimSpace next assigns a cell phenotype *c*_*i*_ ∈ {1, …, *K*} to each candidate location, where *K* is the number of cell phenotypes. This second MRF operates at a *finer biological scale*: it allows both cell-type composition and local cell– cell interaction patterns to vary across tissue niches. Specifically, we model the phenotype configuration 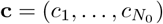 as

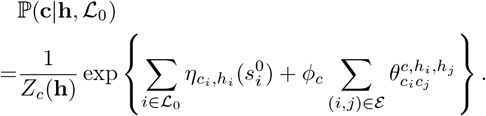

Here, *Z*_*c*_(**h**) is the normalizing constant, the term 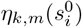 controls the baseline abundance of phenotype *k* in niche *m*, and *ϕ*_*c*_ ≥ 0 controls the overall strength of cell-type spatial dependence. The tensor

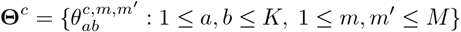

encodes niche-dependent phenotype affinities: 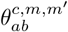 represents the preference for neighboring cells of phenotypes *a* and *b* when their corresponding niches are *m* and *m*′. Positive values promote local co-occurrence of the phenotype pair, whereas negative values discourage adjacency.

Because the neighborhood graph is undirected, the pairwise potential is required to be invariant to the ordering of an edge. We therefore impose 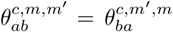. In practice, SimSpace can use a parsimonious niche-specific parameterization: users specify one symmetric phenotype-affinity matrix 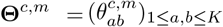 for each niche and set 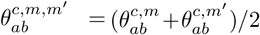 [66]. This parameterization preserves the symmetry required by the undirected MRF while allowing local cell-type interactions to depend on the surrounding niche environment.

The Gibbs update for cell phenotype *c*_*i*_ is

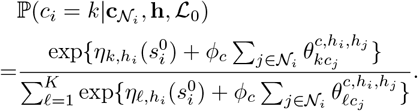

This novel two-level construction separates tissue-scale spatial organization, represented by the niche labels **h**, from local cell-type organization, represented by the phenotype configuration **c**. It therefore enables reference-free generation of spatially coherent niches with distinct, biologically interpretable cell–cell interaction patterns.

#### Cell density and spatial coordinates

To introduce phenotype-dependent cell density, SimSpace applies a thinning step after phenotype assignment. For each phenotype *k*, the user specifies a density parameter *d*_*k*_ ∈ [0, 1], interpreted as the expected retention probability for cells of type *k*. Conditional on the simulated phenotype labels, each candidate location is retained independently according to

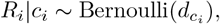

where ℐ= {*i* : *R*_*i*_ = 1} denotes the retained set of cells and *N* = | ℐ| is the final number of simulated cells. This step allows SimSpace to generate phenotype-specific enrichment or depletion while preserving the spatial organization induced by the MRFs.

The retained lattice coordinates are then perturbed to obtain continuous cell locations: 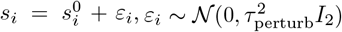 for *i* ∈ ℐ, where *τ*_perturb_ is userspecified. This jittering step converts lattice positions into continuous coordinates and reduces visually apparent grid artifacts. It is not intended to model hard-core cell packing or explicit cell diameters. The spatial dependence in SimSpace is instead generated by the niche-level and cell-phenotype MRFs through the neighborhood graph and affinity parameters. Since the perturbations are independent across cells and independent of the simulated labels, they do not create the spatial correlations evaluated in our benchmarks. Consistently, the spatial summary statistics were stable across a broad range of *τ*_perturb_ values, indicating that the observed spatial structure is driven by the MRF model rather than by coordinate jittering (**Supp Fig. 3C**).

#### Omics feature simulation

After the spatial locations, niches, and cell phenotypes are generated, SimSpace simulates the omics profile

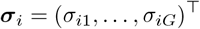

for each retained cell *i* ∈ ℐ, where *G* is the number of synthetic features. These features may represent genes, proteins, chromatin features, or other molecular measurements. Users may either call existing reference-free single-cell simulators, such as Splatter [39], or use the built-in probabilistic simulator described below.

SimSpace also includes a built-in probabilistic model for de novo omics-feature simulation [67]. Throughout, Gamma(*α, β*) denotes a Gamma distribution with shape *α* and scale *β*. First, SimSpace divides *G* synthetic features (genes, proteins, etc.) into universal features and marker features. Universal features represent background molecular programs that are expressed across cell types. For a universal feature *g*, its baseline mean is sampled as

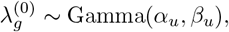

and the count for cell *i* is generated as

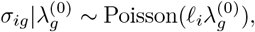

where *ℓ*_*i*_ is an optional cell-specific size factor. If library-size variation is not considered, *ℓ*_*i*_ is set to one.

Marker features encode phenotype-specific molecular signals. Each marker feature *g* is assigned a target phenotype *κ*(*g*) ∈ {1, …, *K*}. We draw a background mean and a marker-effect mean: 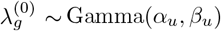 and 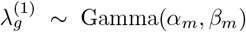 and define the cell-specific mean

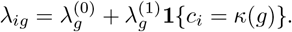

The marker-feature count is then sampled as

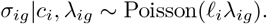

The parameters (*α*_*u*_, *β*_*u*_) and (*α*_*m*_, *β*_*m*_) control the background expression level and the strength of phenotype-specific marker signals, respectively. More overdispersed count distributions can be obtained by retaining the Gamma–Poisson mixture interpretation or by replacing the Poisson sampling step with a negative-binomial model.

#### Spatial ligand–receptor effects

To introduce spatially mediated molecular interactions, SimSpace can designate a set of feature pairs 𝒫_LR_ = {(*g*_*l*_, *g*_*r*_)} as ligand–receptor pairs. For each pair (*g*_*l*_, *g*_*r*_), the expression mean of each feature is multiplicatively modulated by the local neighborhood expression of its partner.

SimSpace implements this mechanism as a *two-stage procedure*. First, baseline means 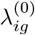 and baseline counts 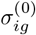 are generated for all features using the universal and marker-feature models described above. For ligand–receptor features, these baseline counts are used to compute local neighborhood-average partner expression. SimSpace then updates the corresponding feature means and resamples the ligand– receptor features from the adjusted means.

Specifically, let 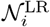 denote the neighborhood of retained cell *i* used for ligand–receptor modulation, and assume 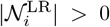. For feature *g*, define the neighborhood-average and global-average baseline expression as

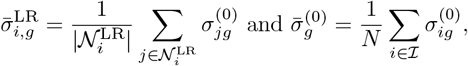

where ℐ is the set of retained cells and *N* = | ℐ |. For a ligand–receptor pair (*g*_*l*_, *g*_*r*_), the adjusted means are

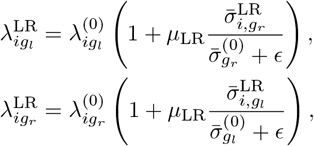

where *µ*_LR_ ≥ 0 controls the strength of the spatial interaction effect and *ϵ* > 0 is a small constant used for numerical stability. The final ligand–receptor feature counts are then generated from the adjusted means, for example

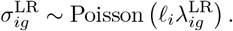

The final simulated count for feature *g* is 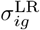 if *g* belongs to a designated ligand–receptor pair, and 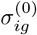 otherwise.

This construction induces spatially localized molecular dependence beyond the cell-phenotype labels alone. For example, cells located near neighborhoods with high receptor expression may exhibit enhanced ligand expression, and vice versa. The parameter *µ*_LR_ provides a direct tuning mechanism for the strength of this spatial molecular interaction.

#### Summary

Overall, the reference-free mode of SimSpace integrates three levels of controlled variation: tissue-scale niche organization, local cell-phenotype interaction, and molecular-feature generation. Unlike simulation schemes that generate spatial coordinates, cell labels, and molecular profiles as largely separate components, SimSpace couples these layers through a shared hierarchical generative model. This design makes the ground-truth spatial architecture, cell-type composition, local interaction structure, marker-feature signals, and ligand–receptor effects simultaneously known and user-controllable. These properties make SimSpace particularly suitable for benchmarking spatial omics methods whose performance depends jointly on spatial organization and molecular variation.

### Parameter sensitivity and tuning

To guide parameter selection in reference-free simulation, we performed a systematic sensitivity analysis of the main parameters controlling spatial structure: the global smoothness parameters of the MRFs, the neighborhood structure, the coordinate-perturbation scale *τ*_perturb_, and the affinity matrices governing niche and cell-phenotype interactions. These parameters control distinct and interpretable aspects of the simulator. The MRF smoothness parameters determine the overall strength of local spatial dependence; the neighborhood graph determines the spatial range over which local interactions operate; *τ*_perturb_ controls the amount of post-sampling coordinate jitter; and the affinity matrices directly encode co-localization or repulsion between niches or cell phenotypes.

We evaluated the effect of these parameters using complementary spatial summary statistics, including Moran’s *I*, Geary’s *C*, Ripley’s centered *L*-function, neighborhood entropy, and cell-type interaction scores (**Supp. Fig. 3**). Across a broad range of settings, the parameters produced stable and biologically interpretable changes in the simulated spatial patterns. Increasing the MRF smoothness parameter strengthened local spatial autocorrelation and generated more spatially coherent domains. Changing the neighborhood structure primarily altered the spatial scale of local interactions, with limited effect on global composition. Varying *τ*_perturb_ affected only the degree of coordinate jitter and did not materially change spatial autocorrelation, indicating that the observed spatial structure is driven by the specified or calibrated MRF model rather than by coordinate jittering. Finally, targeted changes to the entries of the affinity matrices produced predictable changes in niche organization and cell-type co-localization, illustrating how SimSpace can encode user-specified biological hypotheses about local tissue architecture (**Fig. 6C**). Together, these analyses provide empirically grounded tuning guidelines for adapting SimSpace to different tissue types, spatial resolutions, and biological scenarios.

### Spatial summary statistics

To evaluate spatial patterns of the simulated data, we used several spatial summary statistics, which are computed based on the spatial coordinates and cell types of the provided dataset. Let *s*_*i*_ = (*x*_*i*_, *y*_*i*_) denote the spatial coordinate of cell *i*, let *c*_*i*_ 1, …, *K* denote its phenotype label, and let *N* denote the total number of simulated cells. For each cell *i*, let 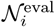 denote its evaluation neighborhood. We define the local neighborhood composition vector ***p***_*i*_ = (*p*_*i*1_, …, *p*_*iK*_)^⊤^, where

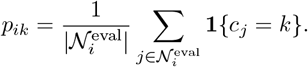

#### Moran’s *I* and Geary’s *C*

We construct a spatial weight matrix *W*_eval_ = (*w*_*ij*_), where *w*_*ij*_ ≥ 0 measures the spatial proximity between cells *i* and *j*. Unless otherwise specified, *W*_eval_ is defined by a *K*_nn_-nearest-neighbor graph with *K*_nn_ = 20, the default choice in SimSpace that reflects a local spatial neighborhood commonly used in spatial transcriptomics studies [12, 68, 69]. Specifically, *w*_*ij*_ = 1 if cell *j* is among the nearest neighbors of cell *i* and *w*_*ij*_ = 0 otherwise. We set *w*_*ii*_ = 0 and write 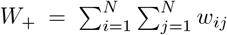. This evaluation graph is used only to compute spatial statistics and need not be identical to the neighborhood graph used in the generative MRF model.

Moran’s *I* and Geary’s *C* quantify spatial autocorrelation in cell-type labels. For each phenotype *k*, define the binary indicator *u*_*ik*_ = **1**{*c*_*i*_ = *k*} and 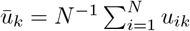. The phenotype-specific Moran’s statistic is

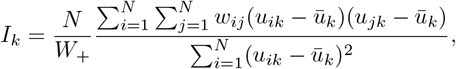

and the corresponding Geary’s statistic is

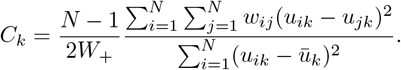

Large positive values of Moran’s *I* indicate that cells of the same phenotype tend to be spatially clustered, whereas larger values of Geary’s *C* indicate greater local dissimilarity. These statistics are computed only for phenotypes with nonzero empirical variance in the simulated sample.

#### Ripley’s centered *L*-function

Ripley’s *K*-function characterizes spatial clustering over a continuum of distance scales. For phenotype *k*, let ℐ_*k*_ = {*i* : *c*_*i*_ = *k*} with *N*_*k*_ = | ℐ_*k*_ |, and let |Ω| denote the area of the spatial domain after coordinate normalization. The phenotype-specific empirical *K*-function is computed as

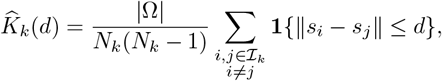

for cell types with *N*_*k*_ ≥ 2. We then transform it as

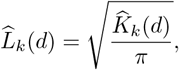

and report the centered statistic 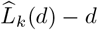, which is approximately centered around zero under complete spatial randomness. Positive values indicate same-type clustering at distance scale *d*, whereas negative values indicate spatial inhibition or dispersion. In our analyses, 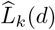 − *d* is evaluated on a fixed grid of distance values after coordinate normalization, yielding one spatial-scale profile per replicate. We summarize this vector within each replicate and visualize the distribution of replicate-level summaries in **Fig. 2D**. For irregular spatial domains, edge correction may be applied to reduce boundary bias. In our benchmark analyses, the same spatial domain and evaluation procedure were used across methods and replicates, so the reported Ripley’s statistics were used primarily for relative comparison.

#### Cell-type interaction score

To summarize local co-localization and repulsion between cell phenotypes, we computed a cell-type interaction matrix from neighborhood compositions, following the principle used in scFeatures [45]. Specifically, the empirical local co-occurrence matrix is 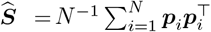. The (*a, b*) entry of 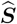 summarizes the frequency with which phenotypes *a* and *b* co-occur in local neighborhoods. To distinguish enrichment from changes in global abundance, we also compare 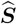 with a permutation null obtained by randomly permuting phenotype labels over fixed spatial coordinates. This yields an interaction score that captures phenotype-pair enrichment or depletion beyond what is expected from marginal cell-type frequencies alone.

#### Neighborhood entropy

Neighborhood entropy measures the local diversity of cell phenotypes. Using the same evaluation neighborhood 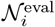, the entropy around cell *i* is defined as 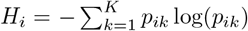, with the convention that 0 log 0 = 0. Larger values indicate more diverse local neighborhoods, whereas smaller values indicate locally homogeneous tissue regions. When comparisons across different numbers of cell types are needed, we use the normalized entropy *H*_*i*_/ log *K*.

To summarize the sample-level distribution of local diversity, we compute the empirical density of *H*_*i*_ over a fixed set of bins 0 = *q*_0_ *< q*_1_ < · · · *< q*_*J*_. Specifically, 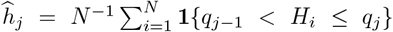 for *j* = 1, …, *J*. The resulting entropy distribution provides a sample-level summary of whether the simulated tissue is dominated by homogeneous domains, highly mixed neighborhoods, or a combination of both.

### Reference-based simulation

SimSpace also supports reference-based simulation, in which spatial parameters are calibrated from an annotated spatial omics dataset. Let

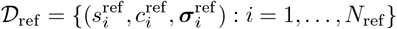

denote the reference dataset, where 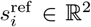 is the spatial coordinate, 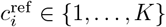 is the annotated cell phenotype, and 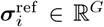 is the omics profile.

The goal is to learn a set of interpretable spatial parameters that reproduce key spatial properties of the reference tissue, while preserving the modular generative structure of SimSpace.

Let ***ϑ*** denote the collection of spatial simulation parameters, including niche-level MRF parameters, cell-phenotype MRF parameters, phenotype-specific density parameters, and coordinate-perturbation parameters when applicable. For a given ***ϑ***, SimSpace generates a synthetic phenotype configuration

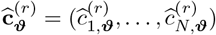

for *r* = 1, …, *R* independent simulation replicates. We calibrate ***ϑ*** by matching spatial summary statistics between the simulated and reference phenotype configurations. Specifically, we define

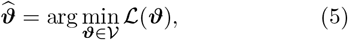

where 𝒱 is the user-specified parameter space and

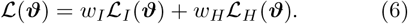

Here, ℒ_*I*_ and ℒ_*H*_ measure discrepancies in spatial autocorrelation and neighborhood-entropy distribution, respectively. For example,

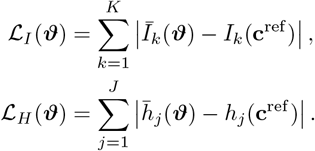

In these expressions, *I*_*k*_(·) is Moran’s *I* for phenotype *k* and *h*_*j*_() is the empirical mass of the neighborhood-entropy distribution in bin *j*. The quantities 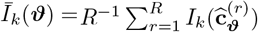 and 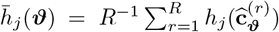 average these statistics over independent synthetic replicates, thereby reducing Monte Carlo variability in the objective. The weights *w*_*I*_, *w*_*H*_ allow users to emphasize different aspects of tissue structure, such as local spatial autocorrelation, neighborhood diversity, or global cell-type composition.

Because the simulator is stochastic and the objective is generally non-smooth, SimSpace uses a genetic algorithm to solve (5). At each generation, candidate parameter vectors are evaluated by running the reference-free simulator and computing the calibration loss in (6). Selection, crossover, and mutation steps then update the candidate population until a prespecified stopping criterion or maximum number of generations is reached. This procedure provides a practical likelihood-free calibration strategy for matching complex spatial phenotypes without requiring closed-form likelihood evaluation for the MRF normalizing constants.

After obtaining 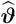, SimSpace generates new spatial patterns by running the calibrated niche-level and cell-phenotype MRFs, followed by the density and coordinate-perturbation steps described above. This produces synthetic tissues that retain the interpretable parameterization of the reference-free model while matching key spatial features of the reference sample.

#### Reference-based omics-profile simulation

Conditional on the simulated coordinates and phenotype labels, omics profiles are generated using a reference-based single-cell simulation module. In the current implementation, SimSpace uses scDesign3 [29] to model cell-type-specific omics profiles. scDesign3 estimates empirical feature distributions from the reference dataset, including mean, dispersion, and zero-inflation components when appropriate. Because these distributions are learned from the reference data rather than imposed through an RNA-specific parametric form, the same framework can be applied across modalities, including transcriptomic counts and protein intensity measurements such as CODEX. For each simulated cell, SimSpace samples an omics profile from the fitted cell-type-specific distribution corresponding to its assigned phenotype. The final output is a synthetic spatial omics dataset containing spatial coordinates, cell phenotypes, and molecular profiles.

### Runtime and memory benchmarking

We benchmarked the runtime and memory usage of SimSpace in reference-free mode and compared it with scCube and scMultiSim, two existing simulators that support de novo generation of spatial patterns. Each method was evaluated on datasets ranging from 100 to 40,000 cells, with five independent replicates for each dataset size. In all settings, each simulator generated both spatial coordinates and omics profiles for 220 features. Benchmarks were performed on a single CPU thread on an Apple M2 chip with 32 GB RAM.

For reference-based simulation, SimSpace additionally performs parameter calibration using a genetic algorithm. Each fitness evaluation consists of one reference-free simulation followed by computation of the spatial calibration loss. Thus, the total runtime scales approximately with the product of the population size, the number of generations, and the cost of a single simulation. Because candidate parameter evaluations are independent, this calibration step is naturally parallelizable. In our implementation, using eight CPU threads reduced the calibration time to approximately 10 minutes for 2,500 cells with population size 60 and 20 generations on comparable hardware.

#### Three-dimensional data simulation

SimSpace extends naturally to three-dimensional spatial omics data. In the 3D setting, each cell has coordinate *s*_*i*_ = (*x*_*i*_, *y*_*i*_, *z*_*i*_) ∈ ℝ^3^, and the neighborhood graph is constructed in three-dimensional space, for example using Euclidean distance thresholds or *K*_nn_-nearest-neighbor rules. The niche-level and cell-phenotype MRFs retain the same pairwise form as in the two-dimensional setting, with edge sets now defined over adjacent spatial locations or voxels in 3D. The Gibbs updates, density-thinning step, and omics simulation procedures remain unchanged. This extension allows SimSpace to generate volumetric spatial omics datasets with controllable tissue-domain structure, local cell-cell interactions, and spatially patterned molecular features.

### Spatial clustering and visualization

We used SpatialPCA and BANKSY to assess whether the simulated datasets preserve the spatial and molecular organization observed in the reference data. For each analysis, we jointly analyzed the SimSpace, scCube, and Xenium reference datasets by assembling a common data object containing the omics profile, spatial coordinates, and cell-type annotation for every cell. Unless otherwise required by a specific method, omics features were normalized using the same preprocessing pipeline across datasets, and spatial coordinates were scaled to a common coordinate system before downstream analysis. This joint analysis design ensured that differences in the resulting embeddings and clusters reflected differences between the simulated and reference spatial omics structures, rather than differences in preprocessing.

For SpatialPCA, we selected the top 100 highly variable genes and used default parameters for model fitting. The first two principal components were used to visualize the dominant spatially structured variation.

For BANKSY, we evaluated multiple values of the spatial weighting parameter, *λ* ∈ { 0.2, 0.5, 0.8}, and visualized the resulting embeddings using UMAP. To isolate the contribution of spatial organization from that of omics-profile simulation, we performed a matched-profile control analysis. In this analysis, we kept the simulated spatial coordinates and celltype labels from SimSpace and scCube fixed, but replaced each simulated omics profile by a profile sampled from the Xenium reference data with the same cell type. Specifically, for each simulated cell *i* with phenotype label *c*_*i*_ = *k*, we sampled a reference cell *r*(*i*) uniformly from 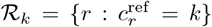, and assigned 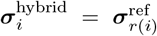. Sampling was performed independently across simulated cells, with replacement when necessary. The resulting hybrid datasets preserve the simulated spatial coordinates and cell-type arrangements, while imposing the same cell-type-specific molecular distributions as the Xenium reference. Therefore, differences observed in SpatialPCA or BANKSY under this control can be attributed primarily to the simulated spatial organization rather than to differences in the omics-profile simulator.

### Benchmarking cell-type deconvolution methods

We used SimSpace-generated datasets to benchmark five cell-type deconvolution methods: cell2location, RCTD, CARD, Seurat, and spatialDWLS. Each method was run following its recommended workflow and default settings unless otherwise stated. For cell2location, the reference model was trained for 3,000 epochs with batch size 256. In the spatial mapping step, we used the default settings *N cells per location = 30* and *detection alpha = 20*, and trained the deconvolution model for 5,000 epochs. For RCTD, we retained all spots and genes by setting *CELL MIN INSTANCE = 1*, and evaluated both the *multi* and *full* modes. For CARD, we set *minCountGene = 1* and *minCountSpot = 1*. For Seurat, we used *SCTransform* to normalize both the reference and spatial datasets, and used the first 30 principal components for anchor-based label transfer. For spatialDWLS, we computed the cell-type mean expression matrix from the Xenium reference data and performed deconvolution with *n cell = 20*.

Performance was evaluated by comparing the estimated and ground-truth cell-type proportions across spatial locations. Let ***A***_*i*_ = (*A*_*i*1_, …, *A*_*iK*_)^⊤^ and ***B***_*i*_ = (*B*_*i*1_, …, *B*_*iK*_)^⊤^ denote the ground-truth and predicted cell-type proportion vectors at spatial location *i*, respectively. We evaluated performance using PCC (computed over all location–cell-type pairs), RMSE, and location-level Jaccard similarity (Jaccard_*i*_ for location *i*), which are computed as:

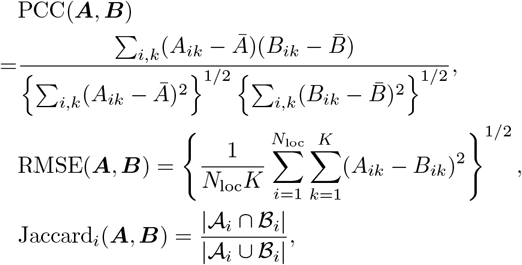

where Ā and 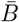 are the corresponding averages over all location–cell-type pairs, *N*_loc_ is the number of evaluated spatial locations, and 𝒜_*i*_ = {*k* : *A*_*ik*_ *>* 0} and ℬ_*i*_ = *k* : *B*_*ik*_ *> ι* with *ι* > 0 being a small threshold for predicted proportions. The final Jaccard score is the average of Jaccard_*i*_(***A, B***) over spatial locations. These metrics assess complementary aspects of deconvolution performance: global agreement in proportion estimates, calibration error, and recovery of the set of present cell types.

### Benchmarking spatially variable gene detection methods

We used SimSpace-generated datasets with known ground-truth SVGs to benchmark six SVG detection methods: Hotspot, SpatialDE, scBSP, SPARK, Giotto, and Moran’s *I* as implemented in Seurat. Each method was run using its recommended pipeline and default parameters unless otherwise specified. For SPARK, we set *percentage = 0* to retain all genes during screening.

Let 𝒢_SVG_ denote the ground-truth set of SVGs and let 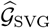 denote the set of genes detected by a given method ^1^. We evaluated each method using rank agreement and binary detection accuracy. For rank-based evaluation, we compared the method-specific gene ranking with the ground-truth ranking induced by the known spatial-effect strength in the simulation. We used Pearson correlation between rank-normalized significance scores and Kendall’s *τ* to quantify ranking agreement.

#### Kendall’s tau

Kendall’s *τ* measures the concordance between two gene rankings via 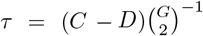, where *C* and *D* are the numbers of concordant and discordant gene pairs, respectively, and *G* is the number of genes. Larger values indicate stronger agreement between the detected ranking and the ground-truth ranking.

#### F1 score

For binary detection, a gene was considered detected if its adjusted significance value was below the prespecified threshold. Given 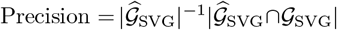 and 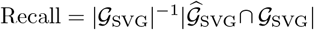, F1 score is defined as

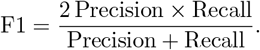

If no genes were detected by a method, precision and F1 score were set to zero. Together, these metrics evaluate whether a method identifies the correct set of SVGs and ranks genes according to their true spatial signal strength.

#### Kendall’s coefficient of concordance (W)

To quantify the stability of gene rankings across independently generated reference-free simulation datasets, we also computed Kendall’s coefficient of concordance, denoted by *W*. Suppose *R* simulation replicates each produce a ranking of the same *G* genes. Let *r*_*rg*_ be the rank of gene *g* in replicate *r*, and let 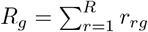 be the rank sum for gene *g*. Kendall’s coefficient of concordance is

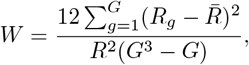

with 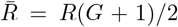. The statistic takes values between 0 and 1, with values near 0 indicating little agreement across replicates and values near 1 indicating highly stable rankings. This analysis assesses whether the simulated spatial signals produce reproducible benchmark rankings across independent datasets.

## Data availability

The Xenium human breast tumor dataset was downloaded from 10X Genomics website: https://www.10xgenomics.com/products/xenium-in-situ/preview-dataset-human-breast. Its cell type annotation can be found in its original publication. The MERFISH mouse hypothalamus dataset was obtained from the original publication and can be downloaded at: https://datadryad.org/dataset/doi:10.5061/dryad.8t8s248. The spatial proteomics dataset used for the CODEX/PhenoCycler-Fusion analyses will be available on the public NYU Langone OMERO server at the time of publication. Code used for image processing, cell segmentation, cell phenotyping, and spatial analysis is available at: https://github.com/FenyoLab/EC_codeximaging

## Code availability

The SimSpace package is available at https://github.com/TianxiaoNYU/simspace as a Python package, where users can find the source code, documentation, and examples for using the package. The code to reproduce all figures in this study is available at https://github.com/TianxiaoNYU/simspace-reproducibility.

## Supplementary Information

**Supplementary Figure 1.**
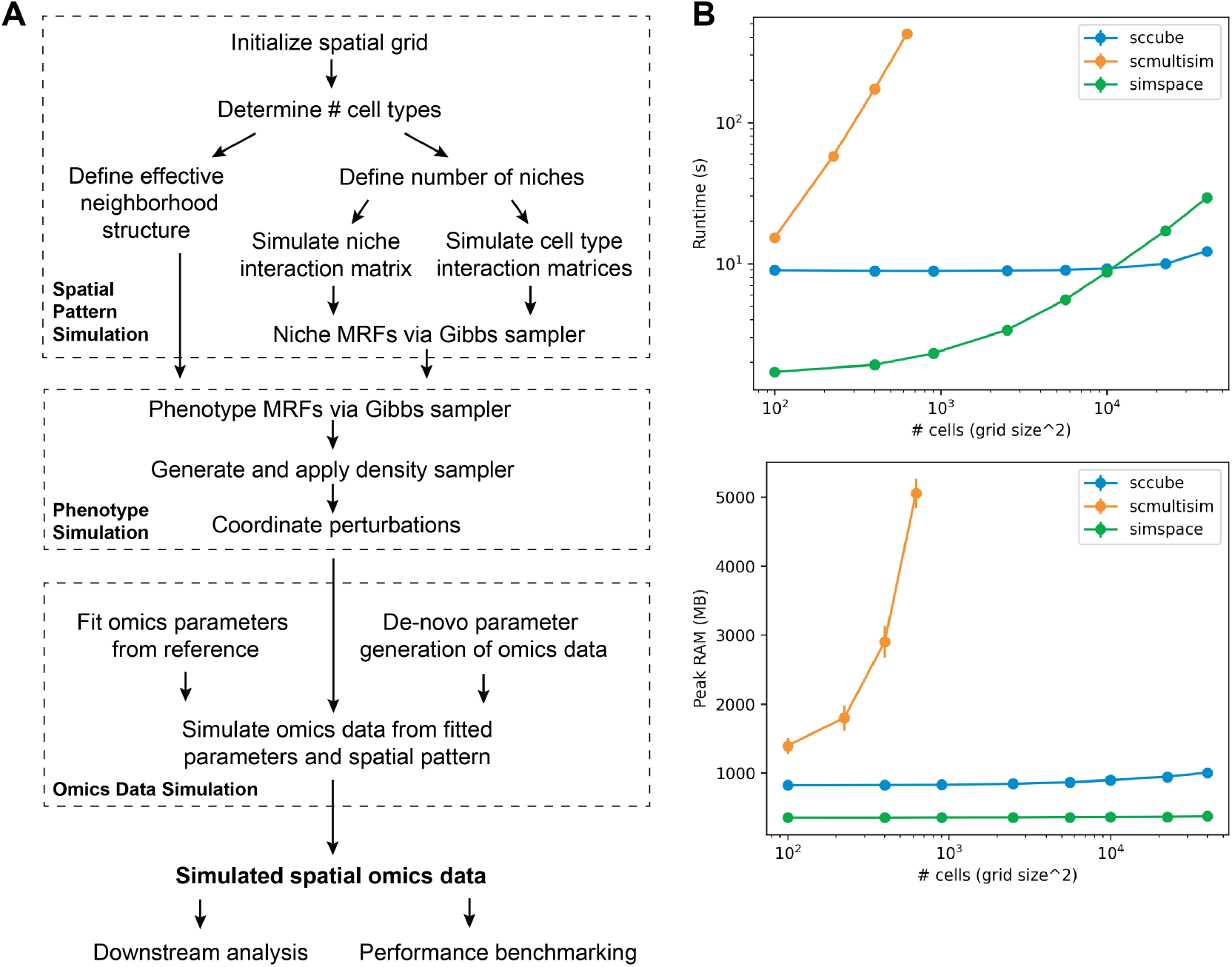
(A) Overview of the SimSpace generative framework. Flowchart summarizing the three main stages of SimSpace simulation: (1) Spatial Pattern Simulation: A spatial grid is initialized, followed by user-defined settings or fitted parameters from reference data for the number of cell types, neighborhood structure, and niche/cell-type affinity. MRFs are simulated via Gibbs sampling to generate structured spatial distributions. (2) Phenotype Simulation: Phenotype-level MRFs are constructed, followed by density sampling and coordinate perturbation to emulate realistic cellular density and variability. (3) Omics Data Simulation: Omics parameters are either fitted from a reference dataset or generated de novo. Simulated spatial patterns are paired with expression profiles to produce final spatial omics datasets. (B) Runtime and memory usage of SimSpace compared with other simulation methods. Benchmarking was performed for SimSpace, scCube, and scMultiSim in reference-free mode, simulating both spatial and omics data for 220 genes across datasets ranging from 100 to 40,000 cells (five replicates each). SimSpace shows competitive runtime and memory efficiency, scaling effectively with increasing dataset size. All methods were run using a single thread on an Apple M2 chip with 32 GB RAM.

**Supplementary Figure 2.**
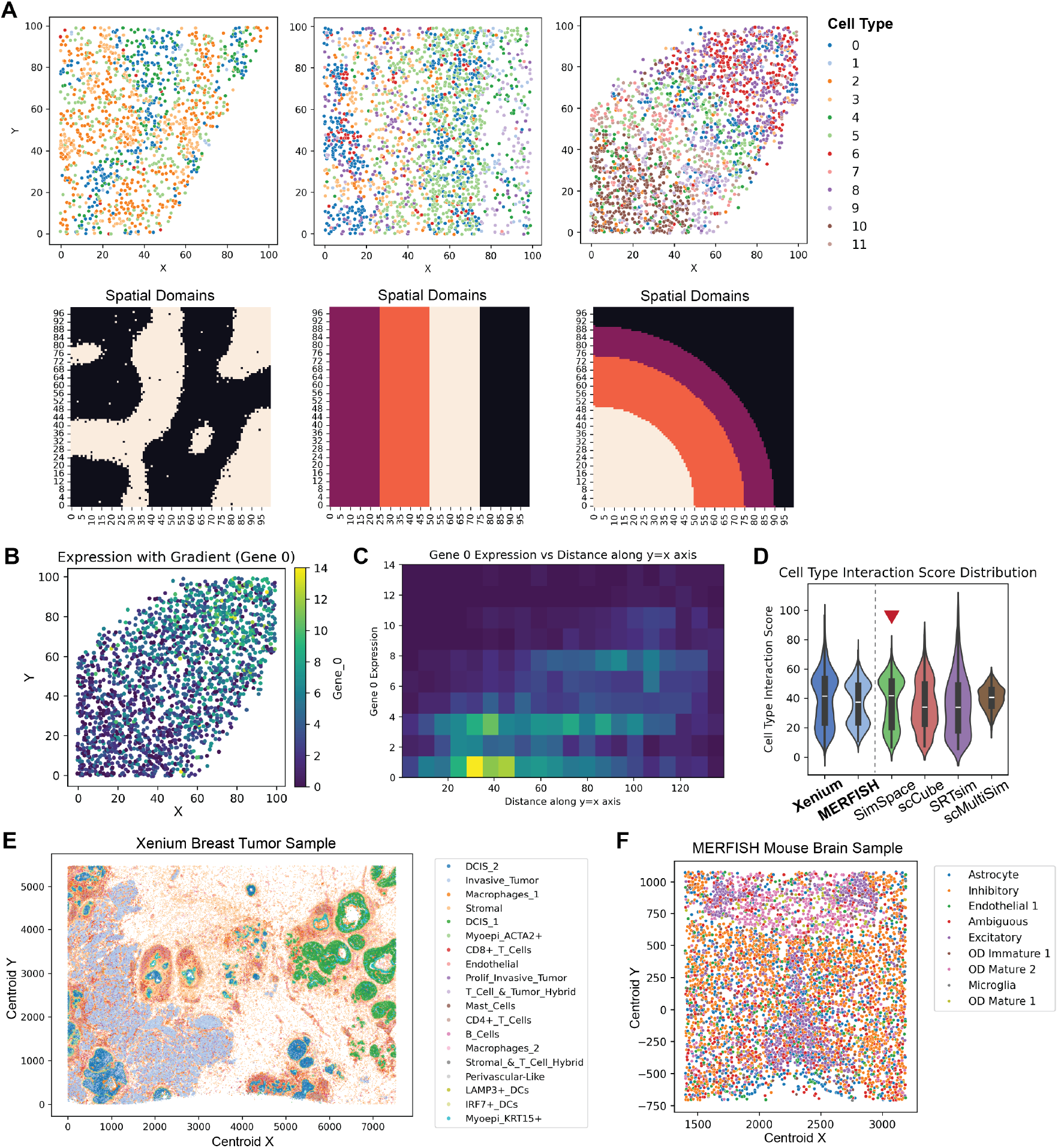
(A) Examples of customized designs for simulation grids and spatial domains. The first row shows spatial maps generated from reference-free simulations using user-defined shapes. The second row visualizes the corresponding spatial domain masks, where different colors denote distinct tissue regions or boundaries. (B) Spatial map of single-cell Gene 0 expression. Each point is a cell positioned by its (x,y) coordinates and colored by Gene 0 expression level, illustrating a diagonal spatial gradient across the tissue. (C) Two-dimensional binned density of Gene 0 expression versus position along the diagonal axis. Heatmap shows the number of cells in each bin as a function of Gene 0 expression (y-axis) and the cell’s projected distance along the y=x axis (x-axis). (D) Comparative analysis of spatial patterns in real and simulated data based on cell-type interaction scores. SimSpace simulations (highlighted with red marks) preserve the characteristic bimodal distributions observed in the Xenium and MERFISH reference datasets more accurately than other reference-free simulation methods, including scCube, SRTsim, and scMultiSim. (E) Visualization of the full-resolution Xenium breast tumor sample colored by cell type labels, highlighting the complex spatial organization of tumor, stromal, and immune populations. Cell annotations are based on the original Xenium dataset publication. (F) Visualization of the MERFISH mouse brain sample colored by cell type labels, demonstrating diverse neuronal and glial populations with distinct spatial distributions. Cell annotations are based on the original MERFISH dataset publication.

**Supplementary Figure 3.**
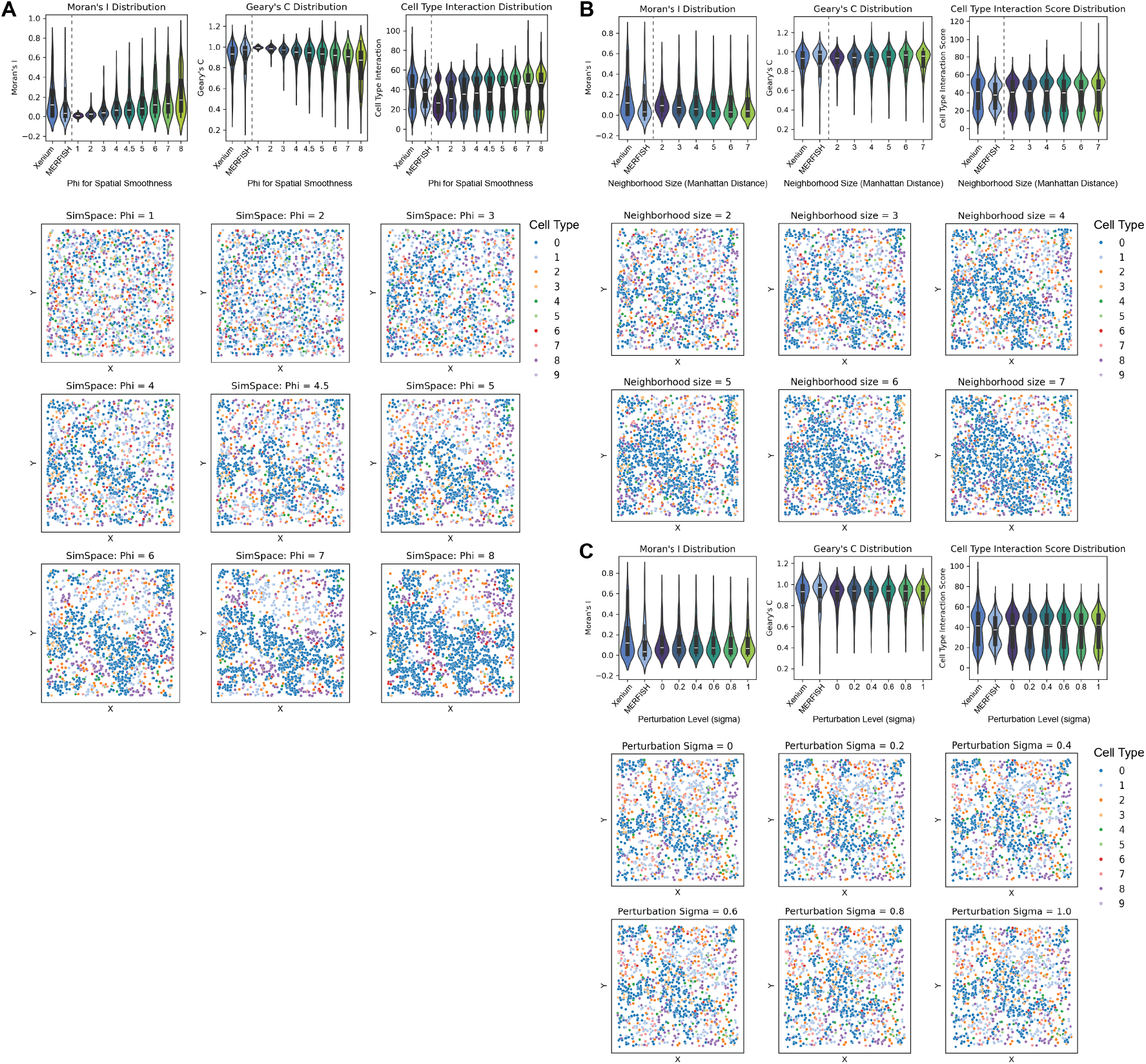
Sensitivity of SimSpace to key spatial modeling parameters. (A) Effect of the spatial smoothness parameter (*ϕ*) on simulated tissues. Increasing *ϕ* strengthens local spatial autocorrelation, as reflected by shifts in Moran’s I, Geary’s C, and cell-cell interaction score distributions. Representative simulated tissues illustrate increasingly coherent spatial niches as *ϕ* increases. (B) Effect of neighborhood size on spatial autocorrelation and interaction patterns. Simulations generated using Manhattan distances of 1-7 exhibit expected changes in Moran’s I, Geary’s C, and interaction scores, with corresponding spatial organization shown below. (C) Effect of coordinate perturbation (*σ*) on spatial structure. As *σ* increases, spatial patterns remain largely intact, with only minor perturbations in cell positions, indicating robustness of the spatial simulation to small coordinate noise.

**Supplementary Figure 4.**
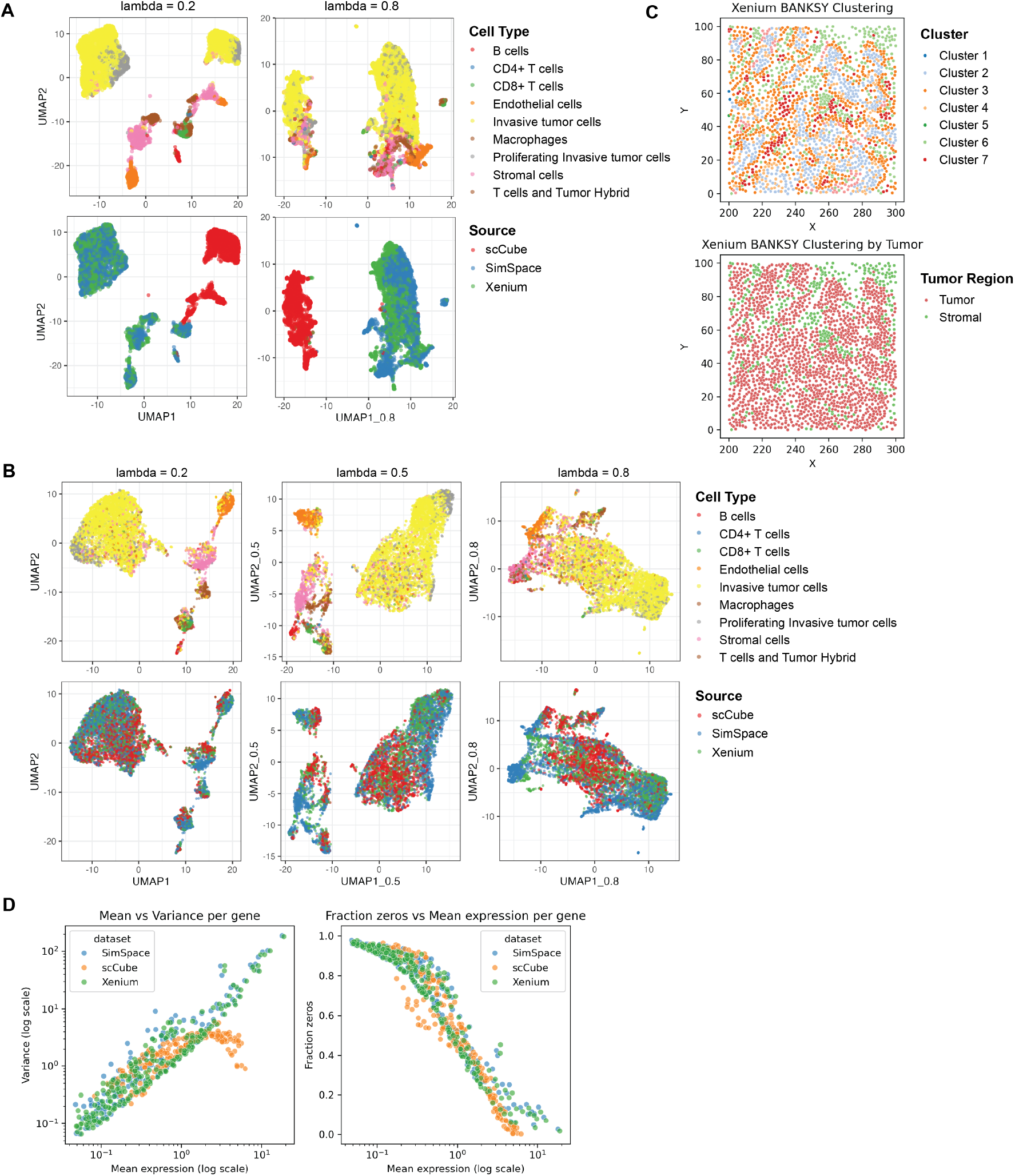
Additional evaluation of reference-based SimSpace simulations. (A) BANKSY UMAP embeddings colored by cell type (top) and data source (bottom). Embeddings were generated using two mixing parameters (*λ* = 0.2 and *λ* = 0.8), which control the contribution of a cell’s own expression versus its neighborhood expression. SimSpace simulations maintain cell-type structure comparable to the Xenium reference while avoiding strong source-driven clustering. (B) BANKSY embeddings computed under different levels of expression-neighborhood mixing (*λ* = 0.2, 0.5, 0.8). When expression profiles are made comparable across datasets, SimSpace continues to produce embeddings that are consistent with the Xenium reference and less biased than those generated by scCube. (C) BANKSY clustering of the Xenium reference tissue. Top: spatial clusters identified by BANKSY across the reference tile. Bottom: clusters annotated by tumor versus stromal domain labels, showing strong agreement between spatially inferred structure and known tissue compartments. (D) Gene-level distributional comparisons between SimSpace simulations, sc-Cube simulations, and Xenium data. Left: mean-variance relationship across genes. Right: fraction of zero counts versus mean expression. SimSpace closely reproduces key gene-level statistical properties of the Xenium reference, while scCube exhibits noticeable deviations, including inflated variance and altered dropout behavior.

**Supplementary Figure 5.**
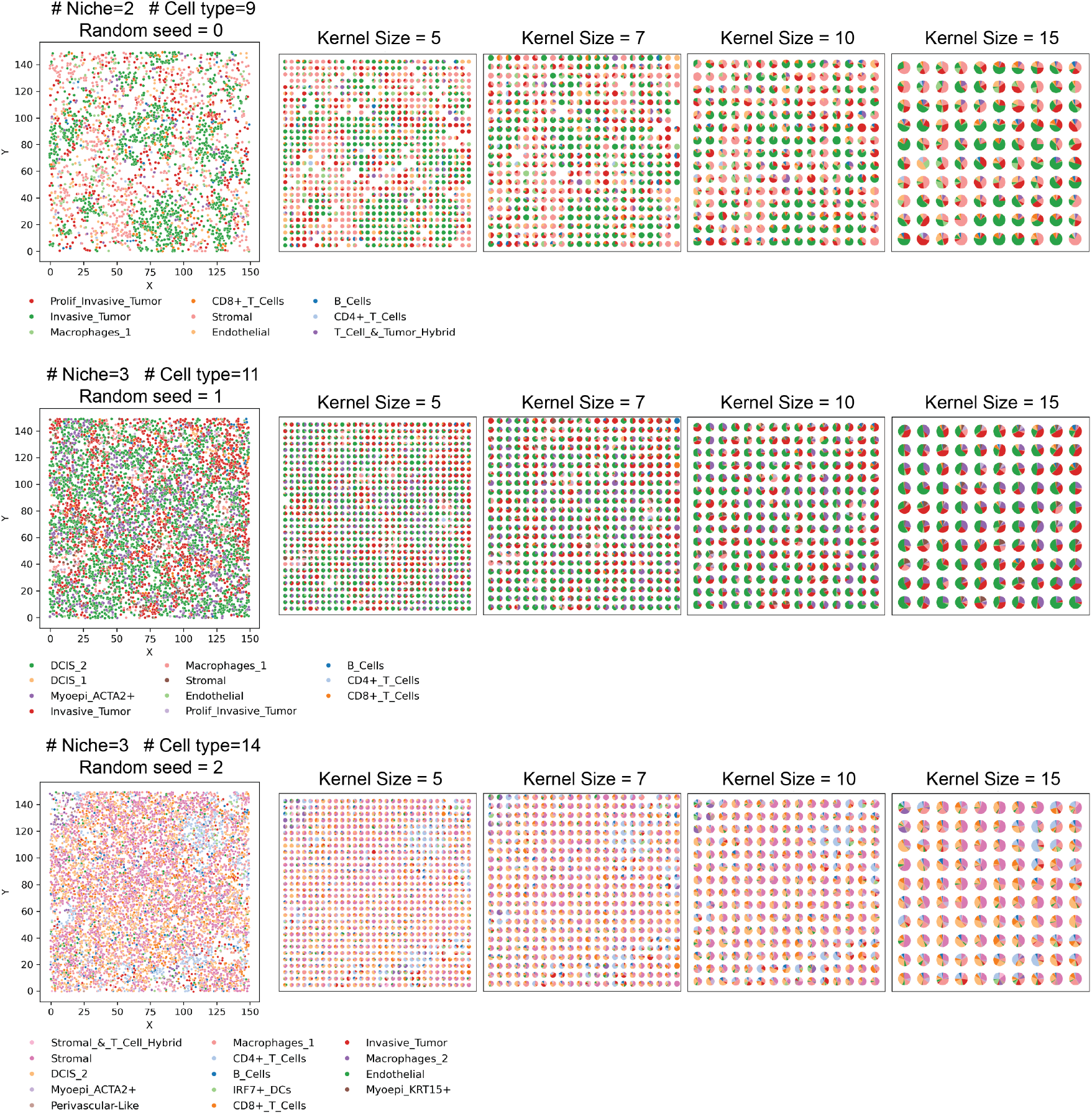
Example SimSpace-generated datasets and their convolved spot-level representations. Three representative simulations derived from Xenium breast tumor tiles are shown, each with different numbers of cell types, spatial niches, and random seeds. For each simulation (rows), the leftmost panel shows the single-cell spatial layout, while the subsequent panels display spot-level data generated by applying manual convolution with kernel sizes of 5, 7, 10, and 15. These examples illustrate how spatial resolution and cellular composition vary across simulated datasets.

**Supplementary Figure 6.**
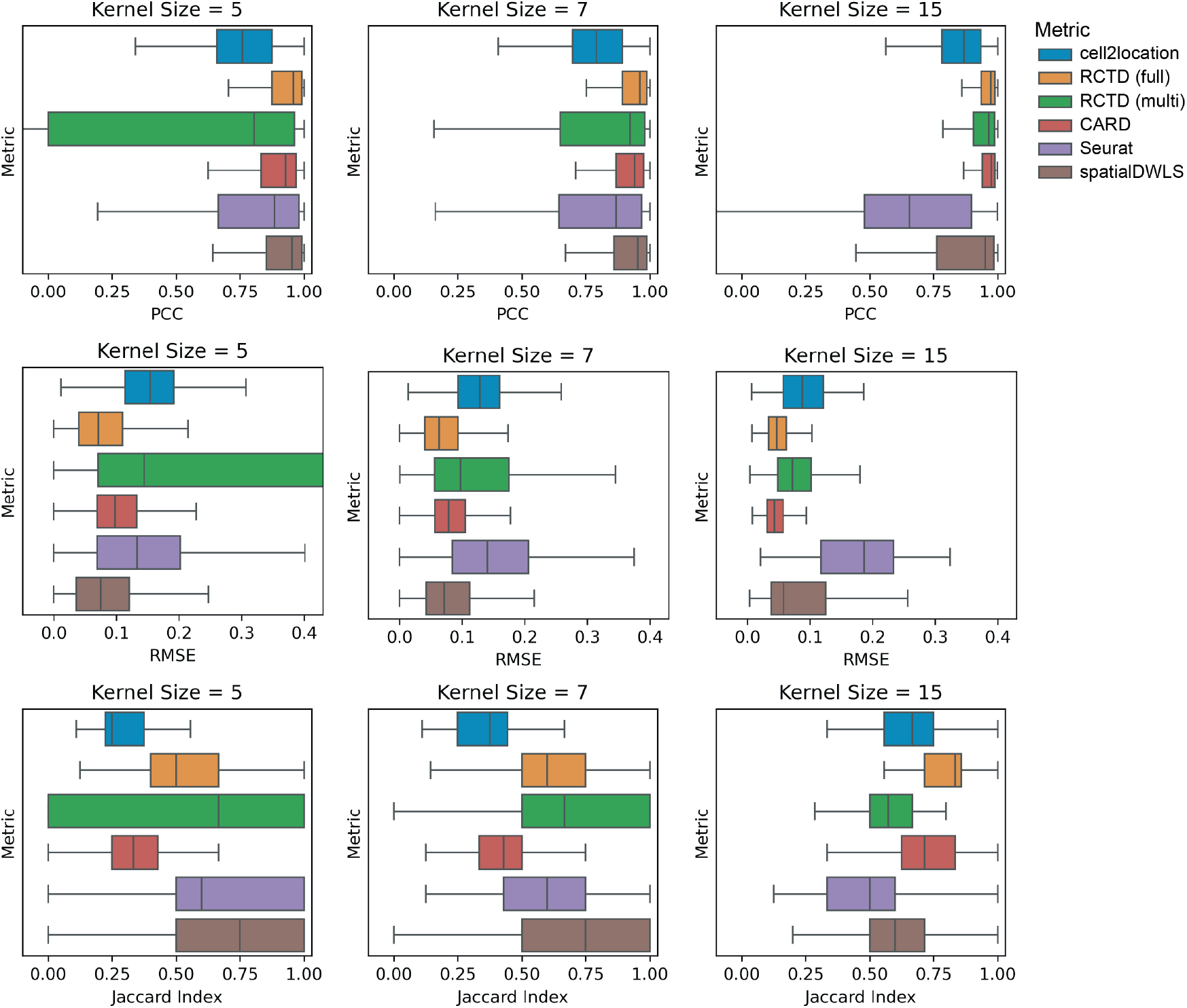
Performance of cell type deconvolution methods across different kernel sizes. Boxplots show benchmarking results for six deconvolution methods (cell2location, RCTD [full and multimode], CARD, Seurat, and spatialDWLS) evaluated on SimSpace-generated datasets. Performance is assessed using PCC (top row), RMSE (middle row), and Jaccard index (bottom row) across three kernel sizes: 5, 7, and 15. Each method’s distribution reflects variation across multiple simulated datasets.

**Supplementary Figure 7.**
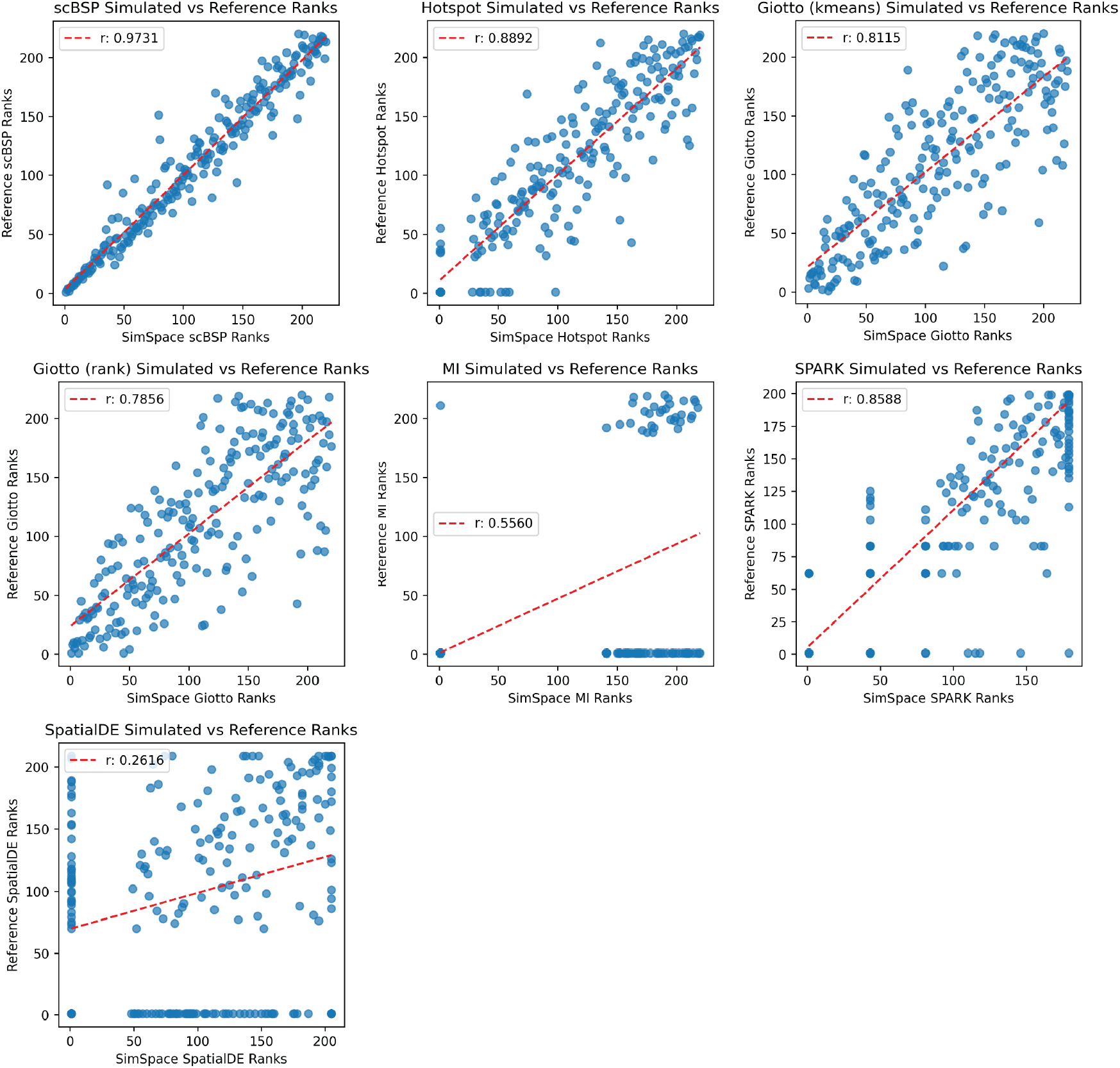
Comparison of SVG rankings between SimSpace simulations and the Xenium reference dataset. Scatter plots compare gene ranks derived from SimSpace reference-based simulations versus the original Xenium reference across seven SVG detection methods: scBSP, Hotspot, Giotto (kmeans), Giotto (rank), Moran’s I, SPARK, and SpatialDE. Each point represents one gene; dotted red lines indicate linear regression fit. Pearson correlation coefficients (r) are shown for each method. scBSP and Hotspot achieve the highest concordance with the reference, while SpatialDE and Moran’s I exhibit lower agreement, suggesting greater sensitivity to spatial variation.

## Footnotes

1 Genes were ranked so that smaller adjusted *p*-values, or larger −log_10_ adjusted *p*-values, corresponded to stronger detected spatial signal.

## References

[1] Z. K. Tuong, K. W. Loudon, B. Berry, N. Richoz, J. Jones, X. Tan, Q. Nguyen, A. George, S. Hori, S. Field, A. G. Lynch, K. Kania, P. Coupland, A. Babbage, R. Grenfell, T. Barrett, A. Y. Warren, V. Gnanapragasam, C. Massie, and M. R. Clatworthy. Resolving the immune landscape of human prostate at a single-cell level in health and cancer. Cell Rep, 37(12):110132, 2021.

[2] L. A. Walsh and D. F. Quail. Decoding the tumor microenvironment with spatial technologies. Nat Immunol, 24(12):1982–1993, 2023.

[3] Z. Yao, C. T. J. van Velthoven, M. Kunst, M. Zhang, D. McMillen, C. Lee, W. Jung, J. Goldy, A. Abdelhak, M. Aitken, K. Baker, P. Baker, E. Barkan, D. Bertagnolli, A. Bhandiwad, C. Bielstein, P. Bishwakarma, J. Campos, D. Carey, T. Casper, A. B. Chakka, R. Chakrabarty, S. Chavan, M. Chen, M. Clark, J. Close, K. Crichton, S. Daniel, P. DiValentin, T. Dolbeare, L. Ellingwood, E. Fiabane, T. Fliss, J. Gee, J. Gerstenberger, A. Glandon, J. Gloe, J. Gould, J. Gray, N. Guilford, J. Guzman, D. Hirschstein, W. Ho, M. Hooper, M. Huang, M. Hupp, K. Jin, M. Kroll, K. Lathia, A. Leon, S. Li, B. Long, Z. Madigan, J. Malloy, J. Malone, Z. Maltzer, N. Martin, R. McCue, R. McGinty, N. Mei, J. Melchor, E. Meyerdierks, T. Mollenkopf, S. Moonsman, T. N. Nguyen, S. Otto, T. Pham, C. Rimorin, A. Ruiz, R. Sanchez, L. Sawyer, N. Shapovalova, N. Shepard, C. Slaughterbeck, J. Sulc, M. Tieu, A. Torkelson, H. Tung, N. Valera Cuevas, S. Vance, K. Wadhwani, K. Ward, B. Levi, C. Farrell, R. Young, B. Staats, M. M. Wang, C. L. Thompson, S. Mufti, C. M. Pagan, L. Kruse, N. Dee, S. M. Sunkin, L. Esposito, M. J. Hawrylycz, J. Waters, L. Ng, K. Smith, B. Tasic, X. Zhuang, et al. A high-resolution transcriptomic and spatial atlas of cell types in the whole mouse brain. Nature, 624(7991):317–332, 2023.

[4] A. Lomakin, J. Svedlund, C. Strell, M. Gataric, A. Shmatko, G. Rukhovich, J. S. Park, Y. S. Ju, S. Dentro, V. Kleshchevnikov, V. Vaskivskyi, T. Li, O. A. Bayraktar, S. Pinder, A. L. Richardson, S. Santagata, P. J. Campbell, H. Russnes, M. Gerstung, M. Nilsson, and L. R. Yates. Spatial genomics maps the structure, nature and evolution of cancer clones. Nature, 611(7936):594–602, 2022.

[5] L. Moses and L. Pachter. Museum of spatial transcriptomics. Nat Methods, 19(5):534–546, 2022.

[6] L. Liu, A. Chen, Y. Li, J. Mulder, H. Heyn, and X. Xu. Spatiotemporal omics for biology and medicine. Cell, 187(17):4488–4519, 2024.

[7] V. Kleshchevnikov, A. Shmatko, E. Dann, A. Aivazidis, H. W. King, T. Li, R. Elmentaite, A. Lomakin, V. Kedlian, A. Gayoso, M. S. Jain, J. S. Park, L. Ramona, E. Tuck, A. Arutyunyan, R. Vento-Tormo, M. Gerstung, L. James, O. Stegle, and O. A. Bayraktar. Cell2location maps fine-grained cell types in spatial transcriptomics. Nat Biotechnol, 40(5):661–671, 2022.

[8] D. M. Cable, E. Murray, L. S. Zou, A. Goeva, E. Z. Macosko, F. Chen, and R. A. Irizarry. Robust decomposition of cell type mixtures in spatial transcriptomics. Nat Biotechnol, 40(4):517–526, 2022.

[9] Y. Ma and X. Zhou. Spatially informed cell-type deconvolution for spatial transcriptomics. Nat Biotechnol, 40(9):1349–1359, 2022.

[10] T. Stuart, A. Butler, P. Hoffman, C. Hafemeister, E. Papalexi, 3rd Mauck, W. M., Y. Hao, M. Stoeckius, P. Smibert, and R. Satija. Comprehensive integration of single-cell data. Cell, 177(7):1888–1902 e21, 2019.

[11] R. Dong and G. C. Yuan. Spatialdwls: accurate deconvolution of spatial transcriptomic data. Genome Biol, 22(1):145, 2021.

[12] D. DeTomaso and N. Yosef. Hotspot identifies informative gene modules across modalities of single-cell genomics. Cell Syst, 12(5):446–456 e9, 2021.

[13] V. Svensson, S. A. Teichmann, and O. Stegle. Spatialde: identification of spatially variable genes. Nat Methods, 15(5):343–346, 2018.

[14] J. Wang, J. Li, S. T. Kramer, L. Su, Y. Chang, C. Xu, M. T. Eadon, K. Kiryluk, Q. Ma, and D. Xu. Dimension-agnostic and granularity-based spatially variable gene identification using bsp. Nat Commun, 14(1):7367, 2023.

[15] Jinpu Li, Mauminah Raina, Yiqing Wang, Chunhui Xu, Li Su, Qi Guo, Ricardo Melo Ferreira, Michael T Eadon, Qin Ma, Juexin Wang, and Dong Xu. scbsp: A fast and accurate tool for identifying spatially variable features from high-resolution spatial omics data. bioRxiv, 2025.

[16] S. Sun, J. Zhu, and X. Zhou. Statistical analysis of spatial expression patterns for spatially resolved transcriptomic studies. Nat Methods, 17(2):193–200, 2020.

[17] R. Dries, Q. Zhu, R. Dong, C. L. Eng, H. Li, K. Liu, Y. Fu, T. Zhao, A. Sarkar, F. Bao, R. E. George, N. Pierson, L. Cai, and G. C. Yuan. Giotto: a toolbox for integrative analysis and visualization of spatial expression data. Genome Biol, 22(1):78, 2021.

[18] N. Moriel, E. Senel, N. Friedman, N. Rajewsky, N. Karaiskos, and M. Nitzan. Novosparc: flexible spatial reconstruction of single-cell gene expression with optimal transport. Nat Protoc, 16(9):4177–4200, 2021.

[19] Manik Kuchroo, Danielle F. Miyagishima, Holly R. Steach, Abhinav Godavarthi, Yutaka Takeo, Phan Q. Duy, Tanyeri Barak, E. Zeynep Erson-Omay, Scott Youlten, Ketu Mishra-Gorur, Jennifer Moliterno, Declan McGuone, Murat Günel, and Smita Krishnaswamy. sparc recovers human glioma spatial signaling networks with graph filtering. bioRxiv, page 2022.08.24.505139, 2022.

[20] J. Hu, X. Li, K. Coleman, A. Schroeder, N. Ma, D. J. Irwin, E. B. Lee, R. T. Shinohara, and M. Li. Spagcn: Integrating gene expression, spatial location and histology to identify spatial domains and spatially variable genes by graph convolutional network. Nat Methods, 18(11):1342–1351, 2021.

[21] V. Singhal, N. Chou, J. Lee, Y. Yue, J. Liu, W. K. Chock, L. Lin, Y. C. Chang, E. M. L. Teo, J. Aow, H. K. Lee, K. H. Chen, and S. Prabhakar. Banksy unifies cell typing and tissue domain segmentation for scalable spatial omics data analysis. Nat Genet, 56(3):431–441, 2024.

[22] L. Shang and X. Zhou. Spatially aware dimension reduction for spatial transcriptomics. Nat Commun, 13(1):7203, 2022.

[23] T. Vu, S. Seal, T. Ghosh, M. Ahmadian, J. Wrobel, and D. Ghosh. Funspace: A functional and spatial analytic approach to cell imaging data using entropy measures. PLoS Comput Biol, 19(9):e1011490, 2023.

[24] A. Sampath Kumar, L. Tian, A. Bolondi, A. A. Hernandez, R. Stickels, H. Kretzmer, E. Murray, L. Wittler, M. Walther, G. Barakat, L. Haut, Y. Elkabetz, E. Z. Macosko, L. Guignard, F. Chen, and A. Meissner. Spatiotemporal transcriptomic maps of whole mouse embryos at the onset of organogenesis. Nat Genet, 55(7):1176–1185, 2023.

[25] G. Yan, S. H. Hua, and J. J. Li. Categorization of 34 computational methods to detect spatially variable genes from spatially resolved transcriptomics data. Nat Commun, 16(1):1141, 2025.

[26] W. Wang, S. Zheng, S. C. Shin, J. C. Chavez-Fuentes, and G. C. Yuan. Ontrac characterizes spatially continuous variations of tissue microenvironment through niche trajectory analysis. Genome Biol, 26(1):117, 2025.

[27] F. W. Townes and B. E. Engelhardt. Nonnegative spatial factorization applied to spatial genomics. Nat Methods, 20(2):229–238, 2023.

[28] L. Shang, P. Wu, and X. Zhou. Statistical identification of cell type-specific spatially variable genes in spatial transcriptomics. Nat Commun, 16(1):1059, 2025.

[29] D. Song, Q. Wang, G. Yan, T. Liu, T. Sun, and J. J. Li. scdesign3 generates realistic in silico data for multimodal single-cell and spatial omics. Nat Biotechnol, 42(2):247–252, 2024.

[30] J. Zhu, L. Shang, and X. Zhou. Srtsim: spatial pattern preserving simulations for spatially resolved transcriptomics. Genome Biol, 24(1):39, 2023.

[31] J. Qian, H. Bao, X. Shao, Y. Fang, J. Liao, Z. Chen, C. Li, W. Guo, Y. Hu, A. Li, Y. Yao, X. Fan, and Y. Cheng. Simulating multiple variability in spatially resolved transcriptomics with sccube. Nat Commun, 15(1):5021, 2024.

[32] H. Li, Z. Zhang, M. Squires, X. Chen, and X. Zhang. scmultisim: simulation of single-cell multi-omics and spatial data guided by gene regulatory networks and cell-cell interactions. Nat Methods, 2025.

[33] A. Janesick, R. Shelansky, A. D. Gottscho, F. Wagner, S. R. Williams, M. Rouault, G. Beliakoff, C. A. Morrison, M. F. Oliveira, J. T. Sicherman, A. Kohlway, J. Abousoud, T. Y. Drennon, S. H. Mohabbat, Teams x Development, and S. E. B. Taylor. High resolution mapping of the tumor microenvironment using integrated single-cell, spatial and in situ analysis. Nat Commun, 14(1):8353, 2023.

[34] J. R. Moffitt, D. Bambah-Mukku, S. W. Eichhorn, E. Vaughn, K. Shekhar, J. D. Perez, N. D. Rubinstein, J. Hao, A. Regev, C. Dulac, and X. Zhuang. Molecular, spatial, and functional single-cell profiling of the hypothalamic preoptic region. Science, 362(6416), 2018.

[35] G. Gut, M. D. Herrmann, and L. Pelkmans. Multiplexed protein maps link subcellular organization to cellular states. Science, 361(6401), 2018.

[36] P Clifford and JM Hammersley. Markov fields on finite graphs and lattices, 1971.

[37] G. R. Cross and A. K. Jain. Markov random field texture models. IEEE Trans Pattern Anal Mach Intell, 5(1):25–39, 1983.

[38] Victor Freguglia and Nancy Lopes Garcia. Inference tools for markov random fields on lattices: The r package mrf2d. Journal of Statistical Software, 101(8):1–36, 2022.

[39] L. Zappia, B. Phipson, and A. Oshlack. Splatter: simulation of single-cell rna sequencing data. Genome Biol, 18(1):174, 2017.

[40] Z. Li, T. Wang, P. Liu, and Y. Huang. Spatialdm for rapid identification of spatially co-expressed ligand-receptor and revealing cell-cell communication patterns. Nat Commun, 14(1):3995, 2023.

[41] Z. Cang, Y. Zhao, A. A. Almet, A. Stabell, R. Ramos, M. V. Plikus, S. X. Atwood, and Q. Nie. Screening cell-cell communication in spatial transcriptomics via collective optimal transport. Nat Methods, 20(2):218–228, 2023.

[42] P. A. P. Moran. Notes on continuous stochastic phenomena. Biometrika, 37(1/2):17–23, 1950.

[43] R. C. Geary. The contiguity ratio and statistical mapping. The Incorporated Statistician, 5(3):115–146, 1954.

[44] B. D. Ripley. The second-order analysis of stationary point processes. Journal of Applied Probability, 13(2):255–266, 1976.

[45] Y. Cao, Y. Lin, E. Patrick, P. Yang, and J. Y. H. Yang. scfeatures: multi-view representations of single-cell and spatial data for disease outcome prediction. Bioinformatics, 38(20):4745–4753, 2022.

[46] X. Liang, M. Torkel, Y. Cao, and J. Y. H. Yang. Multi-task benchmarking of spatially resolved gene expression simulation models. Genome Biol, 26(1):57, 2025.

[47] H. Xu, H. Fu, Y. Long, K. S. Ang, R. Sethi, K. Chong, M. Li, R. Uddamvathanak, H. K. Lee, J. Ling, A. Chen, L. Shao, L. Liu, and J. Chen. Unsupervised spatially embedded deep representation of spatial transcriptomics. Genome Med, 16(1):12, 2024.

[48] P. L. Stahl, F. Salmen, S. Vickovic, A. Lundmark, J. F. Navarro, J. Magnusson, S. Giacomello, M. Asp, J. O. Westholm, M. Huss, A. Mollbrink, S. Linnarsson, S. Codeluppi, A. Borg, F. Ponten, P. I. Costea, P. Sahlen, J. Mulder, O. Bergmann, J. Lundeberg, and J. Frisen. Visualization and analysis of gene expression in tissue sections by spatial transcriptomics. Science, 353(6294):78–82, 2016.

[49] S. G. Rodriques, R. R. Stickels, A. Goeva, C. A. Martin, E. Murray, C. R. Vanderburg, J. Welch, L. M. Chen, F. Chen, and E. Z. Macosko. Slide-seq: A scalable technology for measuring genome-wide expression at high spatial resolution. Science, 363(6434):1463–1467, 2019.

[50] L. Yan and X. Sun. Benchmarking and integration of methods for deconvoluting spatial transcriptomic data. Bioinformatics, 39(1), 2023.

[51] H. Li, J. Zhou, Z. Li, S. Chen, X. Liao, B. Zhang, R. Zhang, Y. Wang, S. Sun, and X. Gao. A comprehensive benchmarking with practical guidelines for cellular deconvolution of spatial transcriptomics. Nat Commun, 14(1):1548, 2023.

[52] G. Yan, S. H. Hua, and J. J. Li. Categorization of 34 computational methods to detect spatially variable genes from spatially resolved transcriptomics data. Nat Commun, 16(1):1141, 2025.

[53] C. Chen, H. J. Kim, and P. Yang. Evaluating spatially variable gene detection methods for spatial transcriptomics data. Genome Biol, 25(1):18, 2024.

[54] Y. Hao, S. Hao, E. Andersen-Nissen, 3rd Mauck, W. M., S. Zheng, A. Butler, M. J. Lee, A. J. Wilk, C. Darby, M. Zager, P. Hoffman, M. Stoeckius, E. Papalexi, E. P. Mimitou, J. Jain, A. Srivastava, T. Stuart, L. M. Fleming, B. Yeung, A. J. Rogers, J. M. McElrath, C. A. Blish, R. Gottardo, P. Smibert, and R. Satija. Integrated analysis of multimodal single-cell data. Cell, 184(13):3573–3587 e29, 2021.

[55] D. Philtron, Y. Lyu, Q. Li, and D. Ghosh. Maximum rank reproducibility: A nonparametric approach to assessing reproducibility in replicate experiments. J Am Stat Assoc, 113(523):1028–1039, 2018.

[56] E. Lundberg and G. H. H. Borner. Spatial proteomics: a powerful discovery tool for cell biology. Nat Rev Mol Cell Biol, 20(5):285–302, 2019.

[57] Y. Goltsev, N. Samusik, J. Kennedy-Darling, S. Bhate, M. Hale, G. Vazquez, S. Black, and G. P. Nolan. Deep profiling of mouse splenic architecture with codex multiplexed imaging. Cell, 174(4):968–981 e15, 2018.

[58] Y. Liu, A. Sinjab, J. Min, G. Han, F. Paradiso, Y. Zhang, R. Wang, G. Pei, Y. Dai, Y. Liu, K. S. Cho, E. Dai, A. Basi, J. K. Burks, K. I. Rajapakshe, Y. Chu, J. Jiang, D. Zhang, X. Yan, P. A. Guerrero, A. Serrano, M. Li, T. H. Hwang, A. Futreal, J. A. Ajani, L. M. Solis Soto, A. A. Jazaeri, H. Kadara, A. Maitra, and L. Wang. Conserved spatial subtypes and cellular neighborhoods of cancer-associated fibroblasts revealed by single-cell spatial multi-omics. Cancer Cell, 43(5):905–924 e6, 2025.

[59] C. C. Hughes. Endothelial-stromal interactions in angiogenesis. Curr Opin Hematol, 15(3):204–9, 2008.

[60] M. V. Patel, Z. Shen, M. Rodriguez-Garcia, E. J. Usherwood, L. J. Tafe, and C. R. Wira. Endometrial cancer suppresses cd8+ t cell-mediated cytotoxicity in postmenopausal women. Front Immunol, 12:657326, 2021.

[61] H. Guan, Q. Xiong, J. Xiong, Y. Liu, and W. Zhang. Cd8+ t cell activation in endometrial cancer: prognostic implications and potential for personalized therapy. Front Immunol, 16:1542669, 2025.

[62] S. Krishnamachari and R. Chellappa. Multiresolution gauss-markov random field models for texture segmentation. IEEE Trans Image Process, 6(2):251–67, 1997.

[63] Q. Zhu, S. Shah, R. Dries, L. Cai, and G. C. Yuan. Identification of spatially associated subpopulations by combining scrnaseq and sequential fluorescence in situ hybridization data. Nat Biotechnol, 2018.

[64] Z. Wu, A. E. Trevino, E. Wu, K. Swanson, H. J. Kim, H. B. D’Angio, R. Preska, G. W. Charville, P. D. Dalerba, A. M. Egloff, R. Uppaluri, U. Duvvuri, A. T. Mayer, and J. Zou. Graph deep learning for the characterization of tumour microenvironments from spatial protein profiles in tissue specimens. Nat Biomed Eng, 6(12):1435–1448, 2022.

[65] Daphne Koller and Nir Friedman. Probabilistic Graphical Models: Principles and Techniques - Adaptive Computation and Machine Learning. The MIT Press, 2009.

[66] Jinyuan Chang, Wen Zhou, Wen-Xin Zhou, and Lan Wang. Comparing large covariance matrices under weak conditions on the dependence structure and its application to gene clustering. Biometrics, 73(1):31–41, 2017.

[67] Abhishek Sarkar and Matthew Stephens. Separating measurement and expression models clarifies confusion in single-cell rna sequencing analysis. Nature genetics, 53(6):770–777, 2021.

[68] J. Li, Y. Shyr, and Q. Liu. aknno: single-cell and spatial transcriptomics clustering with an optimized adaptive k-nearest neighbor graph. Genome Biol, 25(1):203, 2024.

[69] R. Jiang, Z. Li, Y. Jia, S. Li, and S. Chen. Sinfonia: Scalable identification of spatially variable genes for deciphering spatial domains. Cells, 12(4), 2023.

